# Regulation of retinal amacrine cell generation by miR-216b and Foxn3

**DOI:** 10.1101/2020.10.28.358069

**Authors:** Huanqing Zhang, Pei Zhuang, Ryan M. Welchko, Manhong Dai, Fan Meng, David L. Turner

## Abstract

The mammalian retina contains a complex mixture of different types of neurons. We find that the microRNA miR-216b is preferentially expressed in postmitotic retinal amacrine cells in the mouse retina, and expression of miR-216a/b and miR-217 in the retina depend in part on Ptf1a, a transcription factor required for amacrine cell differentiation. Surprisingly, ectopic expression of miR-216b, or the related miR-216a, can direct the formation of additional amacrine cells in the developing retina. In addition, we observe the loss of bipolar neurons in the retina after miR-216b expression. We identify the mRNA for the transcriptional regulator Foxn3 as a retinal target of miR-216b by Argonaute PAR-CLIP and reporter analysis. Inhibition of Foxn3 in the postnatal developing retina by RNAi also increases the formation of amacrine cells and reduces bipolar cell formation, while overexpression of Foxn3 inhibits amacrine cell formation prior to the expression of Ptf1a. Disruption of Foxn3 by CRISPR in embryonic retinal explants also reduces amacrine cell formation. Co-expression of Foxn3 can partially reverse the effects of ectopic miR-216b on retinal cell type formation. Our results identify Foxn3 as a novel regulator of interneuron formation in the developing retina and suggest that miR-216b likely regulates expression of Foxn3 and other genes in amacrine cells.

## Introduction

Cellular diversity is a fundamental feature of the nervous system: networks comprised of different types of neurons and glia form neural circuits with distinct functional properties. The mouse retina has six major classes of neurons that can be further divided into over 100 different subtypes (Clark et al., 2019; Macosko et al., 2015; Shekhar et al., 2016; Yan et al., 2020). This complex mixture of cell types performs initial visual processing in the retina. During retinal development in vertebrates, specific cell types are generated in a broadly consistent order, and individual retinal progenitor cells can give rise to complex combinations of differentiated cell types (reviewed in (Cepko, 2014)). Multiple transcriptional regulators and signaling pathways control the potential of retinal progenitor cells to generate specific cell types at different developmental times (Clark et al., 2019; Konstantinides et al., 2015; Mattar et al., 2015).

Amacrine cells are retinal interneurons that participate in local processing of visual information in the retina (Masland, 2012). Over 60 subtypes of amacrine cells have been described, based on neurotransmitter usage, morphology, and gene expression (Yan et al., 2020). During embryonic and early postnatal development, a series of transcription factors have been shown to regulate the formation of amacrine cells, including the Forkhead protein Foxn4, the pancreas associated transcription factor Ptf1a, Neurod1, Neurod4, and Ascl1 basic-helix-loop-helix factors, the Rorb nuclear receptor, the Meis1 homeobox, and the zinc finger transcription factor Plagl1/Zac1 (Fujitani et al., 2006; Inoue et al., 2002; Islam et al., 2013; Li et al., 2004; Liu et al., 2013; Liu et al., 2020; Ma et al., 2007; Nakhai et al., 2007). Several of these factors also regulate the formation of horizontal cells, another class of retinal interneuron, and/or other retinal neurons.

Noncoding RNAs also have been implicated as regulators of retinal development, including microRNAs (miRNAs). Mature miRNAs are ~22nt small RNAs that, when loaded onto Argonaute proteins, function as negative regulators of mRNA stability and translation (reviewed in (Bartel, 2018)). Target-miRNA interactions usually involve base-pairing between the miRNA seed sequence (positions 2-8 of the mature miRNA) and a complementary sequence in the mRNA, most often located in the 3’ untranslated region (3’ UTR) of the mRNA. Loss of mature miRNAs after retinal-specific disruption of the miRNA processing enzyme Dicer leads to the altered production of multiple retinal cell types, including fewer amacrine cells, with prolonged generation of early cell types (Davis et al., 2011; Georgi and Reh, 2010) and/or widespread cell death (Damiani et al., 2008), depending on the specific timing of Dicer disruption. miRNAs have been implicated in the differentiation or maintenance of retinal neurons and Müller glia, as well as in the neurogenic potential of Müller glia (Kara et al., 2019; Reh and Hindges, 2018; Walker and Harland, 2009; Wohl et al., 2019; Wohl et al., 2017; Xiang et al., 2017).

Here we show that ectopic expression of the miR-216b miRNA during mouse retinal development leads to an increase in amacrine cells and a decrease in bipolar cells. miR-216b is normally expressed in differentiated amacrine cells. We identify the 3’ UTR of the mRNA encoding the Foxn3 transcription factor as a target of miR-216b. Inhibition of Foxn3 by RNAi or CRISPR disruption in developing retinas leads to an increase in amacrine cells, while overexpression of Foxn3 reduces amacrine cell number, and co-expression of Foxn3 with miR-216b can partially reverse the effects of miR-216b. Our observations identify Foxn3 as a novel regulator of retinal interneuron formation and suggest that miR-216b is a regulator of gene expression in amacrine cells.

## Results

To identify miRNAs enriched in retinal amacrine cells, we used small RNA sequencing to compare miRNA levels in embryonic day 18.5 (E18.5) retinas isolated either from mice heterozygous or homozygous for disruption of the gene encoding Ptf1a, a transcription factor required for formation of most amacrine and horizontal cells in the retina (Fujitani et al., 2006; Nakhai et al., 2007). Several miRNAs were reduced in the retinas from the Ptf1a knockout, relative to retinas from Ptf1a heterozygotes, with miR-216b-5p the most downregulated miRNA (Fig.1A, Supplemental Table S1). miR-216b is part of a genomic miRNA cluster with miR-216a and miR-217; miR-216a-5p/3p and miR-217-5p also were decreased in the Ptf1a knockout retinas. Real-time quantitative reverse-transcription PCR (qRT-PCR) confirmed downregulation of the predominant mature miRNA from each of these three miRNA genes in Ptf1a knockout retinas relative to retinas from heterozygotes (Fig. 1B). A developmental time course of retinal expression by qRT-PCR showed that all three mature miRNAs decreased from postnatal day 0 (P0) to P7, but expression was maintained in P12 and adult retinas. In contrast, miR-183-5p, an unrelated miRNA expressed in photoreceptors and bipolar cells (Xu et al., 2007; Zhuang et al., 2020), increased between P0 and adult (Fig. 1C). miRNA in situ hybridization for mature miR-216b-5p revealed expression in the inner retina at P0 and in the inner nuclear layer (INL) and in the ganglion cell layer (GCL) at P12, with the strongest signal at P12 in the inner INL (Fig.1D). Cells labeled by miR-216b-5p in situ hybridization overlapped with amacrine cells, identified by immunodetection of the transcription factor AP2α (Tfap2a) (Bassett et al., 2007) at P0 or P12 (Fig 1E, H). miR-216b-5p also overlaps with AP2α at E18.5 in retinas from Ptf1a heterozygotes, but miR-216b-5p was reduced and AP2α was absent in Ptf1a knockout retinas (Supplemental Fig. S1). In situ hybridization for the miR-216b-5p, miR-216a-5p, and miR-217-5p miRNAs in retinas at different ages revealed expression in INL and GCL (Fig. 1D and Supplemental Fig. S2). At P0, the three miRNAs were detected in the inner retina, but were not present in the neuroblast layer. Reduced expression of these miRNAs in Ptf1a mutant retinas and the localization to the inner retina are consistent with expression of all three miRNAs in postmitotic amacrine cells.

**Fig. 1.**
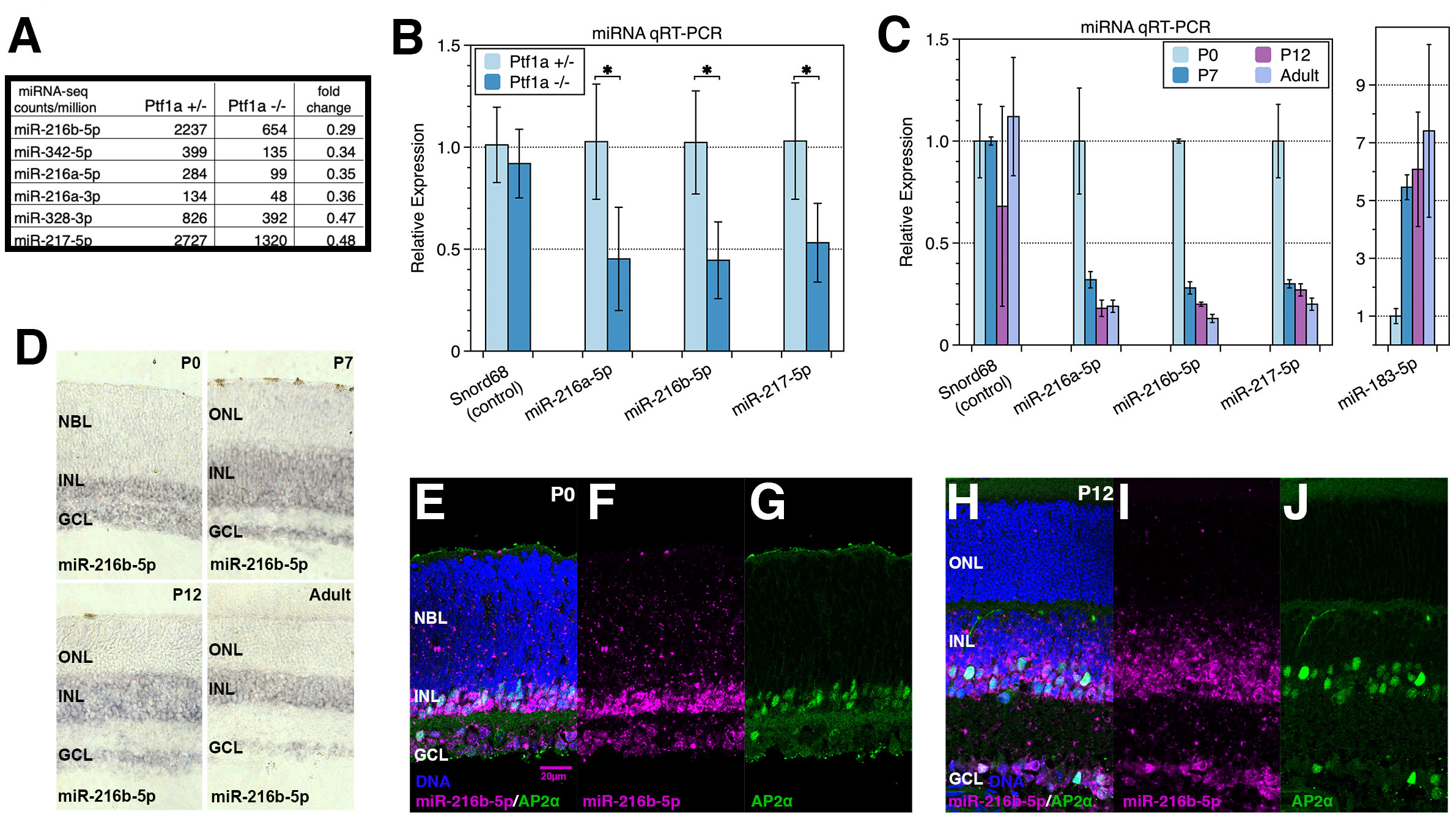
Expression of miR-216a/b and miR-217 miRNAs in the mouse retina. (A) Table of miRNAs with the largest fold decrease in average small RNA sequencing counts in retinas from E16.5 Ptf1a homozygous null mice, relative to retinas from Ptf1a heterozygous mice (among miRNAs with minimum 100 counts per million reads in averaged Ptf1a heterozygous samples; N=2). (B) qRT-PCR for mature miR-216a-5p, miR-216b-5p, and miR-217-5p shows reduced expression of all three miRNAs in E16.5 Ptf1a homozygous retinas, relative to heterozygous retinas (N=3). (C) Mature miR-216a-5p, miR-216b-5p, and miR-217-5p expression decreases in wild-type retinas after P0, but expression persists in adult retina. In contrast, the rod and bipolar miRNA miR-183-5p increases after P0 in the same samples (N=3). (D) miR-216b-5p expression, detected by miRNA in situ hybridization (purple), overlaps with differentiated retinal cells in inner retina at P0 and is restricted to the INL and GCL in the P7, 12, and adult retinas, with the strongest signal in the amacrine cell layer. (E-J) miR-216b-5p expression detected by miRNA fluorescent in situ hybridization (magenta), overlaps with the amacrine cell specific AP2α protein detected by immunofluorescence (green) at P0 (E-G) and P12 (H-J). In this and subsequent figures: data represent mean ± SD, * P < 0.05; NBL: neuroblast layer, INL: inner nuclear layer, GCL: ganglion cell layer, ONL: outer nuclear layer; scale bars as indicated.

The miR-216a-5p and miR-216b-5p miRNAs have similar seed sequences and potentially could bind to overlapping sets of target sites, while miR-217-5p has a distinct seed sequence and would be expected to bind to different target sites (Fig. 2A). To determine whether miR-216a/miR-216b/217 could influence retinal development we ectopically expressed these miRNAs in neonatal mouse retinas. We used DNA plasmid vectors that express GFP with pre-miR-216b, pre-miR-216a, or pre-miR-217 expressed from the intron of the same transcription unit (Fig. 2A), either together or individually, introduced into P0 mouse retinas by *in vivo* electroporation (Matsuda and Cepko, 2004; Zhang et al., 2012). GFP expression was used to identify electroporated cells. A plasmid vector expressing GFP and a pre-miR-155-based shRNA against luciferase was used as a control (a functional synthetic miRNA with no target in the retina (Zhang et al., 2012)). Forced expression of miR-216a, miR-216b, or all three miRNAs together led to a significant increase in the frequency of amacrine cells among GFP-labeled cells at P7, relative to the control, as assessed by immunodetection of AP2α (Fig. 2B-G). Introduction of miR-216b or miR-216a individually yielded similar increases in amacrine cells among the GFP-labeled cells, while introduction of miR-217 by itself did not (Fig. 2E, G). While control retinas had few or no GFP-labeled displaced amacrine cells in the GCL, we observed occasional GFP-labeled amacrine cells in the GCL after electroporation with miR-216b, miR-216a, or all three miRNAs together. In addition to the increase in amacrine cells among the GFP-labeled cells with miR-216a/b, we observed a decrease in GFP-labeled cells in the outer part of the INL (Fig. 2C, D, F), suggesting a decrease in bipolar neurons among the electroporated cells. While antibody markers specific for bipolar cells did not work well at P7, additional experiments described below show that miR-216b does decrease the number of bipolar cells among the electroporated cells when scored at P12.

**Fig. 2.**
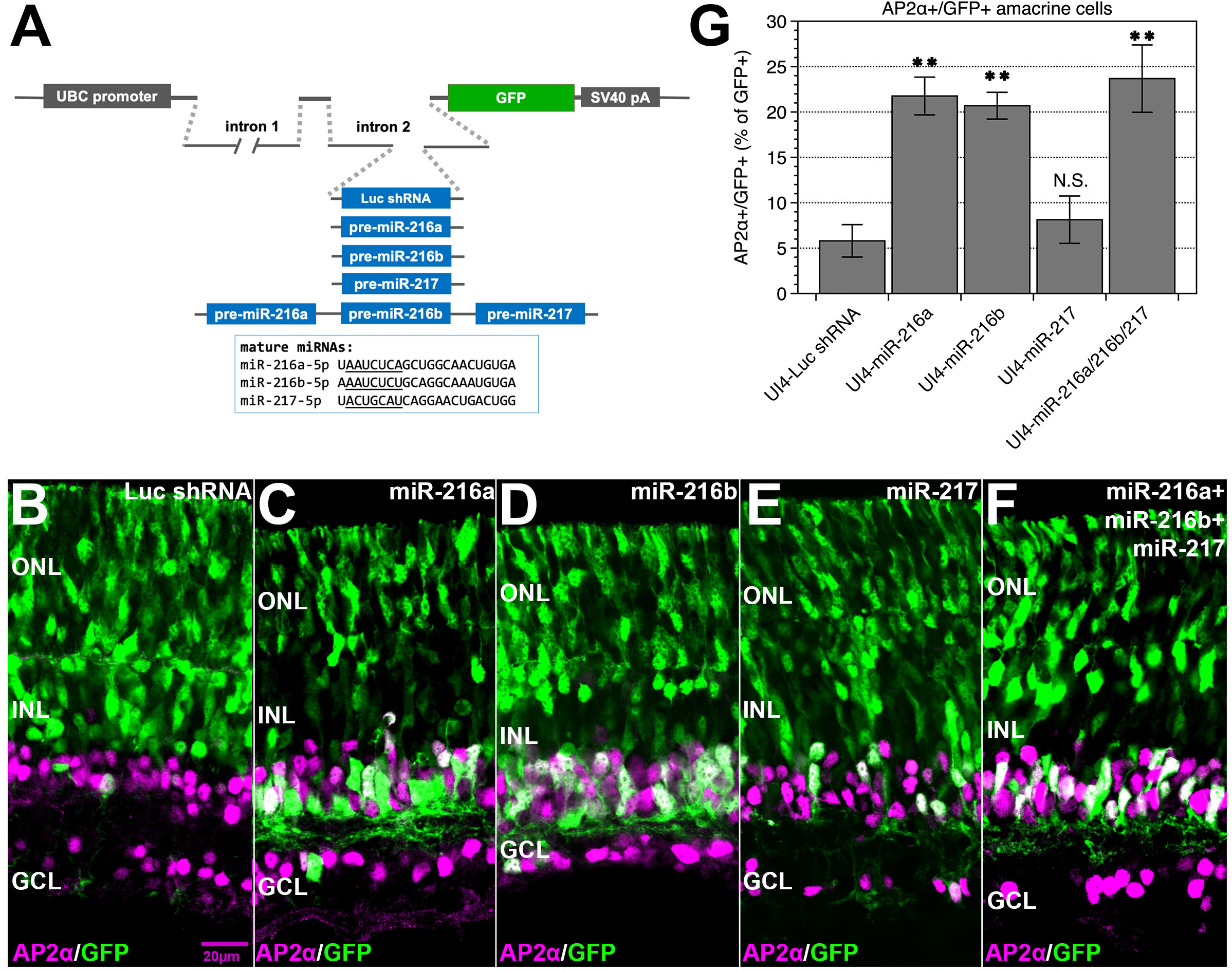
miR-216a/b can promote amacrine cell formation in the developing mouse retina. (A) Schematic of plasmid miRNA expression vectors, which co-express GFP with one or three pre-miRNA(s), or a control Luc shRNA, from an intron within the GFP mRNA. Mature miRNA sequences are shown, with seed sequences underlined. Plasmid vectors were introduced into the retina by *in vivo* electroporation at P0; retinas were analyzed at P7. (B) Widespread GFP+ cells were observed in the ONL and outer INL, with some GFP-labeled AP2α expressing amacrine cells after introduction of the control vector (B). Expression of miR-216a (C) or miR-216b (D), but not miR-217 (E) led to increased AP2α+ amacrine cells among GFP+ cells, and loss of GFP+ cells in the outer part of the INL. Co-expression of all three miRNAs was similar to expression of miR-216a or miR-216b individually (F). Quantitation is shown in (G). ** P < 0.01. N.S.: not significant. Labels as in Fig. 1.

Homozygous disruption of Ptf1a leads to nearly complete loss of AP2α expressing amacrine cells in retinas. Since the expression of miR-216a/b in part depends on Ptf1a, we investigated whether expression of miR-216b would be sufficient to direct the formation of amacrine cells in retinas without Ptf1a. The expression vectors for miR-216b or the Luc shRNA control were introduced by electroporation into E16.5 retinas from mice heterozygous or homozygous for disruption of Ptf1a. Since Ptf1a knockout mice are not viable after birth, retinas were maintained as explants *in vitro* and assessed for AP2α expression at 8 days *in vitro* (DIV). Loss of Ptf1a abolished AP2α-labeled amacrine cells, and the miR-216b expression vector failed to generate AP2α cells in the absence of Ptf1a (Supplemental Fig. S3), indicating that expression of miR-216b cannot direct formation of amacrine cells in the retina without Ptf1a.

We used Argonaute PAR-CLIP (Hafner et al., 2010) to identify potential endogenous miRNA targets in the developing retina, including targets of miR-216a/b. P0 mouse retinas were cultured as explants in 4-thiouridine (4SU) containing media for 1 DIV. After UV crosslinking and lysis, endogenous Argonaute 1-4 proteins with crosslinked RNA were immunoprecipitated and cDNA libraries were prepared from the RNA fragments using a modified PAR-CLIP method, then analyzed by high throughput sequencing (see Materials and Methods for details). We generated five independent Argonaute immunoprecipitated libraries and two IgG immunoprecipitated control libraries. PAR-CLIP with 4SU frequently generates T to C substitutions at the site of crosslinking during reverse transcription, allowing single base resolution of crosslink positions (Hafner et al., 2010). To distinguish crosslinks from random sequencing errors, we identified mapped sequences with consistent T to C substitutions at the same genomic position in two or more of the independent Argonaute PAR-CLIP libraries, without a T to C substitution at the same position in the control libraries. For multiple closely spaced crosslinks, we identified the most frequent crosslink (+/− 15 nt around each crosslink) to create a set of 14,054 most frequent, consistent crosslink sites, which we refer to as predominant crosslinks. Annotation of the predominant crosslink sites revealed that most sites were located in 3’ untranslated regions (UTRs) of mRNAs, with coding regions the next the most frequent location (Fig. 3A), similar to prior PAR-CLIP analyses of Argonaute binding sites (Hafner et al., 2010; Lipchina et al., 2011). As the retina lysates included both cytoplasm and nuclei, nuclear Argonaute likely contributed to the recovery of crosslinked sites in introns and ncRNAs (Sarshad et al., 2018).

**Fig. 3.**
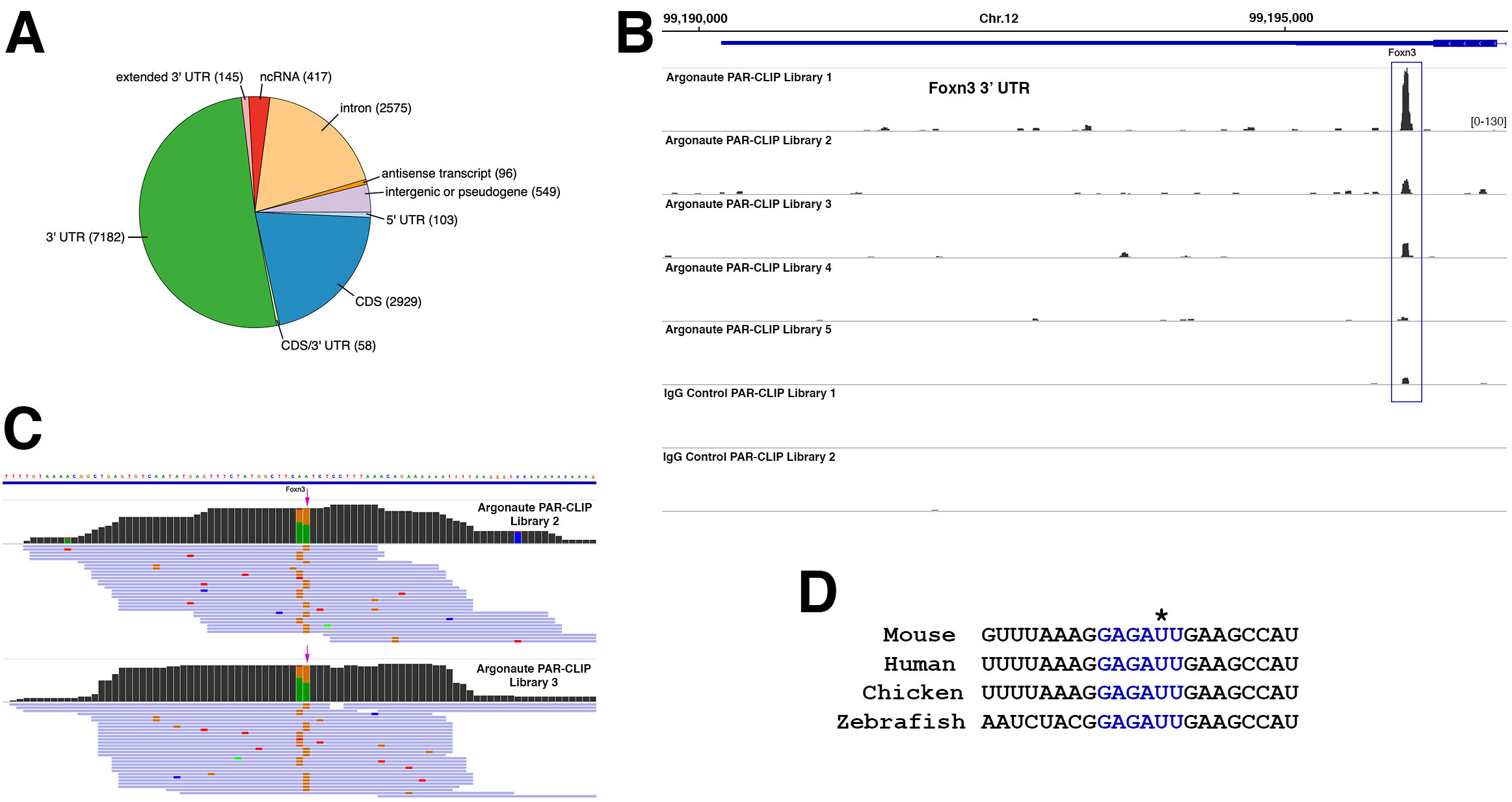
Argonaute PAR-CLIP identifies Foxn3 as a miR-216a/b target. (A) 14,054 predominant Argonaute PAR-CLIP crosslink sites (see text) were mapped to genomic annotations. Most crosslink sites were located in 3’ UTRs, but sites were present in other genomic features (Extended 3’ UTR: <1kb 3’ to an annotated 3’ UTR). (B) A peak of overlapping PAR-CLIP read coverage (blue box) was present in the 3’ UTR of the Foxn3 gene, with reads present in all 5 Argonaute libraries, but not in two IgG control libraries. Same vertical scale used for all libraries. (C) Genomic read mapping in the Foxn3 3’ UTR showing the peak region from Argonaute libraries 2 and 3. Foxn3 is transcribed from the minus strand, so crosslinks appear as A to G substitutions (orange) on the genomic plus strand. The predominant crosslink site is indicated by magenta arrows. (D) The miR-216a/b seed match (blue) present near the predominant crosslink site in the mouse Foxn3 3’ UTR (indicated by *) is conserved in the Foxn3 mRNA in other vertebrates.

Argonaute PAR-CLIP frequently generates crosslinks within a few nucleotides of a bound miRNA seed sequence (Hafner et al., 2010). We looked at the distribution of sequences complementary to miRNA seed sequences within +/− 15nt windows centered on a subset of predominant crosslink sites that mapped to 3’ UTRs. Seed sequence matches for 15 miRNA seeds present in P0 retinas were strongly enriched near crosslink sites. (Supplemental Fig. S4, Supplemental Table 2). Based on these results, and additional results below, we conclude that Argonaute PAR-CLIP identified retinal miRNA binding sites. Here we analyze potential miR-216a/b sites in the Argonaute PAR-CLIP data; analysis of other retinal miRNA binding sites will be described elsewhere.

Seed matches were identified for miR-216a/b-5p near 404 predominant crosslink positions (Supplemental Tables 3, 4). miR-216a-5p and miR-216b-5p differ at position 8 (the last base of the seed, Fig. 2A). Since both miRNAs function similarly when expressed in retinas, we focused on sites with a 6nt match to positions 2-7 of both miRNAs, without a match to a more abundant miRNA seed or a seed match located closer to the crosslink site. We identified a set of 57 predominant crosslink sites that met these criteria (Supplemental Table 3). The site with the largest number of crosslinks near a miR-216a/b-5p 2-7 seed match was located in the 3’ UTR of the Foxn3 transcription factor, and crosslinks at this site were present in all five Argonaute PAR-CLIP libraries (Fig. 3B, C). We also assessed evolutionary conservation of predicted seed sites and the miR-216a/b-5p seed match in the Foxn3 3’ UTR is conserved to chickens and fish (Fig. 3D, Supplemental Table 3). We constructed a Luciferase reporter with a partial 3’ UTR from Foxn3, under the control of a constitutive promoter (Fig. 4A). The Foxn3 3’ UTR fragment included both the frequently crosslinked miR-216a/b-5p site, as well as a second potential miR-216a/b-5p site near the stop codon. The second site had T to C substitutions in two distinct reads in only one retinal PAR-CLIP library and therefore was not identified as a consistent crosslink site. In HEK293 cells in culture, the Foxn3 3’ UTR reporter was inhibited by co-transfection of the miR-216b expression vector but not the miR-216b-mut expression vector, which expresses a version of miR-216b with a seed sequence mutation in miR-216b-5p (and compensating changes in the 3’ end of miR-216b-3p to allow folding of the precursor). A reporter in which the seed matches at both potential sites in the Foxn3 3’ UTR fragment were mutated was not inhibited by the miR-216b vector. However, the miR-216b-mut vector could downregulate the reporter with the mutated miR-216a/b-5p sites, as we used site mutations in the Foxn3 3’ UTR fragment that were complementary to the mutated miR-216b-5p seed sequence in miR-216b-mut (Fig. 4A, B). These observations indicate that the Foxn3 3’ UTR is a miR-216b-5p target.

**Fig. 4.**
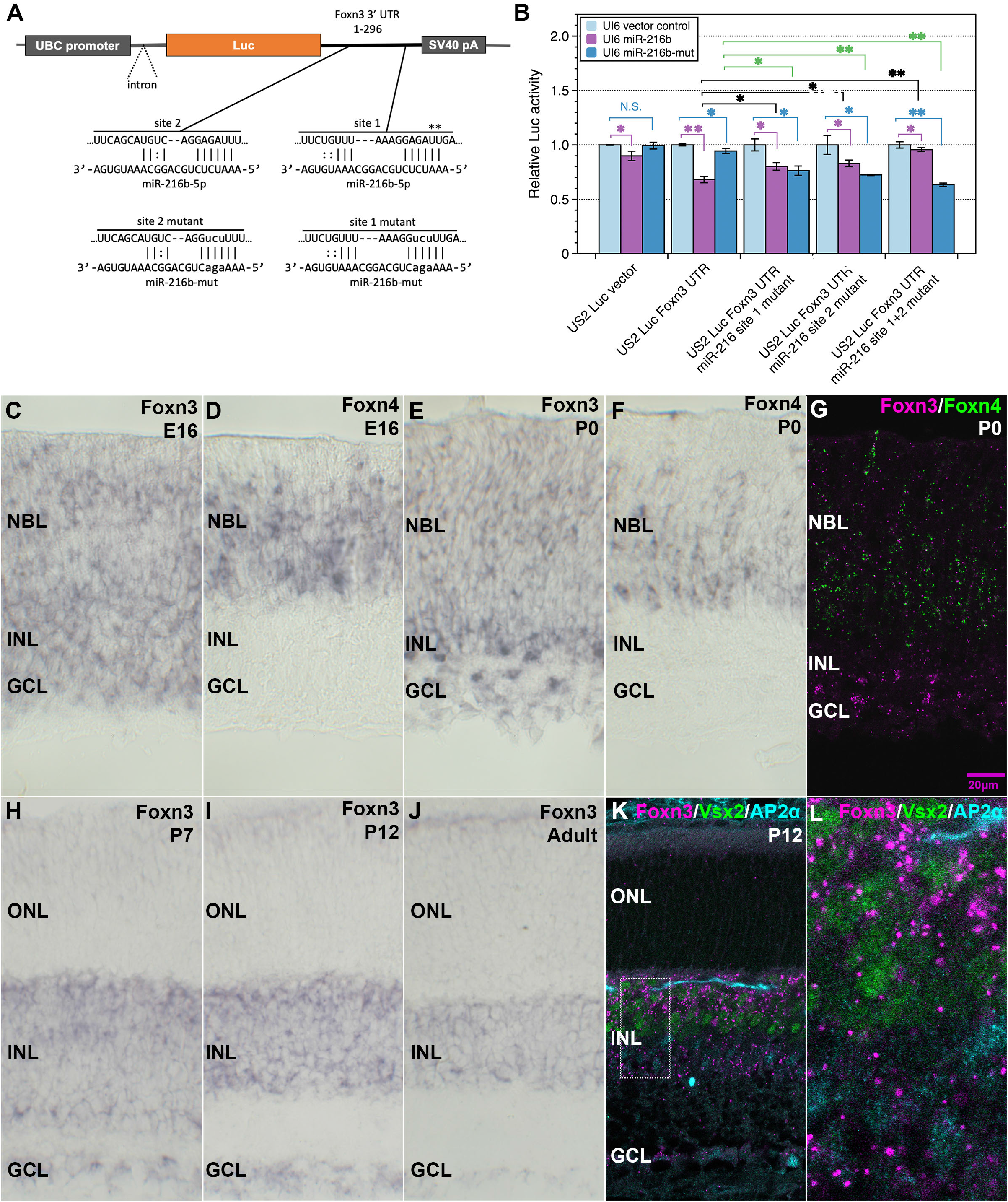
Repression of the 3’ UTR of Foxn3 by miR-216b; Foxn3 mRNA expression during retinal development overlaps with Foxn4 expression. (A) Schematic of a luciferase reporter with 1-296 of the Foxn3 3’ UTR which contains two miR-216a-5p/b-5p seed matches as indicated (most frequent Argonaute PAR-CLIP crosslinking sites for site 1 indicated by asterisks). The miR-216b-mut miRNA, with an altered miR-216b-5p seed sequence is shown, as well as Foxn3 3’ UTR site mutants which it can pair with (lower case indicates substituted bases). (B) The Foxn3 luciferase reporter is repressed by co-expression of miR-216b in P19 cells, but not by co-expression of miR-216b-mut. Mutation of both seed matches in the reporter to match the seed in miR-216b-mut prevents repression by miR-216b and allows repression by miR-216b-mut, while single seed match mutations allow repression by either miRNA. Error bars indicate standard deviation. (C-F, H-J) Expression of Foxn3 and Foxn4 mRNAs (purple) in the developing retina at indicated ages, as detected by in situ hybridization. Foxn3 is expressed throughout development, in the neuroblast layer and in a subset of differentiated cells in the INL/GCL, while Foxn4 is restricted a subset of cells to the neuroblast layer. (G) Overlap between Foxn3 (magenta) and Foxn4 (green) expression in cells of the P0 retina, detected by in situ HCR. (K) Foxn3 mRNA in the INL at P12 (magenta), detected by in situ HCR, overlaps with Vsx2, a marker for bipolar cells (green) as well as a small number of amacrine cells (AP2α, cyan), detected by immunofluorescence. (L) enlargement of boxed region in (K). * P < 0.05, ** P < 0.01. Labels as in Fig. 1.

Intriguingly, Foxn3 is closely related to Foxn4, a transcription factor required for the initial steps of amacrine cell formation (Li et al., 2004). However, while Foxn4 is a transcriptional activator (Lelievre et al., 2012; Liu et al., 2013), Foxn3 is a transcriptional repressor (Scott and Plon, 2005). This difference suggested that Foxn3 might inhibit amacrine cell formation, and that miR-216a/b could inhibit Foxn3 to allow increased amacrine cell formation. Foxn3 is expressed in the head and eye in vertebrates, and loss of Foxn3 function has been shown to lead to a small eye phenotype in Xenopus and in mice, as well as head neural crest defects (Samaan et al., 2010; Schuff et al., 2007). However, a role for Foxn3 in the generation of amacrine cells or other specific retinal cell types has not been described. We therefore further analyzed the expression of Foxn3 during mouse retinal development. At E16.5 or P0, we detected widespread expression of Foxn3 mRNA in the mouse retina by in situ hybridization, both in the neuroblast layer and in some differentiating cells of the inner retina (Fig. 4C, E). We compared Foxn3 expression with Foxn4, which is expressed in retinal precursor cells that can generate amacrine interneurons. Foxn4 mRNA was restricted to cells in the neuroblast layer at E16.5 or P0, consistent with prior reports (Li et al., 2004), and a subset of cells expressed both Foxn3 and Foxn4 mRNAs at P0 or E16.5 (Fig.4C-G, Supplemental Figure 5). In P7, P12, and adult retinas, Foxn3 expression was restricted to the INL and GCL, with most expressing cells located in outer half of the INL (Fig. 4H-J) at P7 and P12. Foxn3 mRNA expression in the INL overlaps with markers of bipolar cells and a small subset of amacrine cells at P12 (Fig.4K, L; Supplemental Fig. S6). Based on recent single cell sequencing data for amacrine cells (Yan et al., 2020), Foxn3 mRNA expression was most correlated with genes expressed in cholinergic amacrine cells, such as Sox2, and the highest level of Foxn3 mRNA was present in cholinergic amacrine cells (Supplemental Table 5).

To determine if Foxn3 could affect amacrine cell formation, we inhibited Foxn3 by RNAi in developing retinas. We used RNA polymerase II driven shRNA vectors that co-express GFP and miR-155-based shRNAs (Chung et al., 2006). We constructed shRNAs against Foxn3 and confirmed their ability to reduce the endogenous Foxn3 mRNA in a mouse cell line using qRT-PCR (Supplemental Fig. S7). The shRNA vectors were delivered into P0 retinas by *in vivo* electroporation. In parallel, we electroporated the control Luc shRNA vector, the expression vector for miR-216b, or the miR-216b-mut expression vector. We observed an increased number of AP2α-positive amacrine cells among the GFP-labeled cells in retinas collected at P12, with two different shRNAs against Foxn3, relative to the control shRNA vector (Fig 5A-C, K). miR-216b expression also increased the fraction of amacrine cells among the GFP-labeled cells at P12, while miR-216b-mut expression did not (Fig 5D, E, K). The fraction of Vsx2 (also known as Chx10) positive bipolar cells among the GFP-labeled cells decreased with either Foxn3 RNAi or miR-216b (Fig. 5F-I, L). Expression of miR-216b-mut led to a smaller decrease in bipolar cell number among GFP-labeled cells, significantly less than the decrease with wild-type miR-216b (Fig 5J, L). These results indicate that the miR-216b vector likely alters retinal development via target repression through the miR-216b-5p strand/seed, and that inhibition of Foxn3 in the developing retina yields changes in retinal cell types similar to changes in cell types seen after overexpression of miR-216b.

**Fig. 5.**
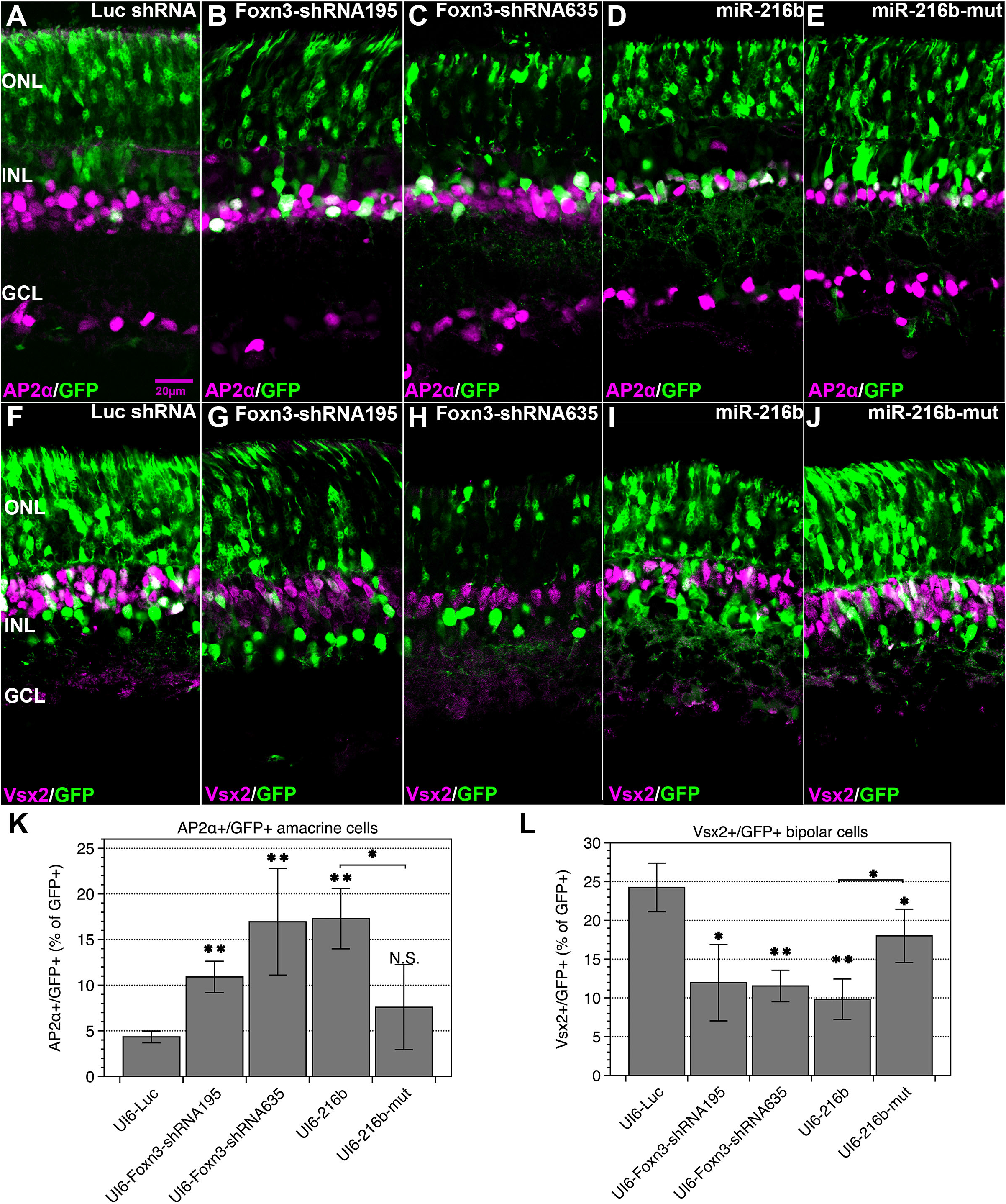
Foxn3 RNAi increases amacrine cells and reduces bipolar cells, while increased amacrine cell formation after miR-216b expression depends on the miR-216b-5p seed sequence. Plasmid vectors that co-express GFP (green) with shRNAs or miRNAs were introduced into the retina by *in vivo* electroporation at P0; retinas were analyzed at P12. (A-C, K) Expression of either of two shRNAs targeting Foxn3 increased GFP-labeled amacrine cells at P12 relative the Luc shRNA control vector (N=3). (D-E, K) The miR-216b vector increased GFP-labeled amacrine cells (AP2α, magenta) at P12 relative to the Luc shRNA, while a vector expressing GFP and miR-216b-mut did not. (F-J, L) Expression of either Foxn3 shRNA or miR-216b reduced GFP-labeled bipolar cells (Vsx2, magenta), while miR-216b-mut reduced GFP-labeled bipolar cells less than miR-216b (N=3). Quantitation is shown in (K,L). * P < 0.05, ** P < 0.01. Labels as in Fig. 1.

As an independent test of Foxn3 function, we used CRISPR/Cas9 to disrupt the Foxn3 gene in embryonic retina explants in culture, then assessed changes in amacrine cell number. Two different Cas9/sgRNA expression vectors, each expressing GFP and two independent sgRNAs against Foxn3, or a control Cas9/GFP vector were electroporated into mouse retinas at E16.5, just before the peak of amacrine cell generation, and the retinas then were maintained as explants for 8 DIV. We observed an increased fraction of AP2α+ cells among the GFP-labeled cells in explants with either of the CRISPR vectors targeting Foxn3, relative to the control vector (Fig. 6A-D). To confirm Cas9 disruption of the Foxn3 gene in retinal cells after electroporation with CRISPR vectors at E16.5, we amplified the four genomic target regions by PCR from GFP positive regions of dissected retina explants at 2 DIV, then detected indels by Illumina sequencing of the PCR products. For each of the vectors targeting Foxn3, one of the two sgRNAs was effective (Supplemental Fig. S8). The increase in amacrine cell fraction after CRISPR disruption of Foxn3 is similar to the results observed after postnatal inhibition of Foxn3 by RNAi. Both of these results indicate that reduced Foxn3 function in cells of the developing retina permits increased amacrine cell formation.

**Fig. 6.**
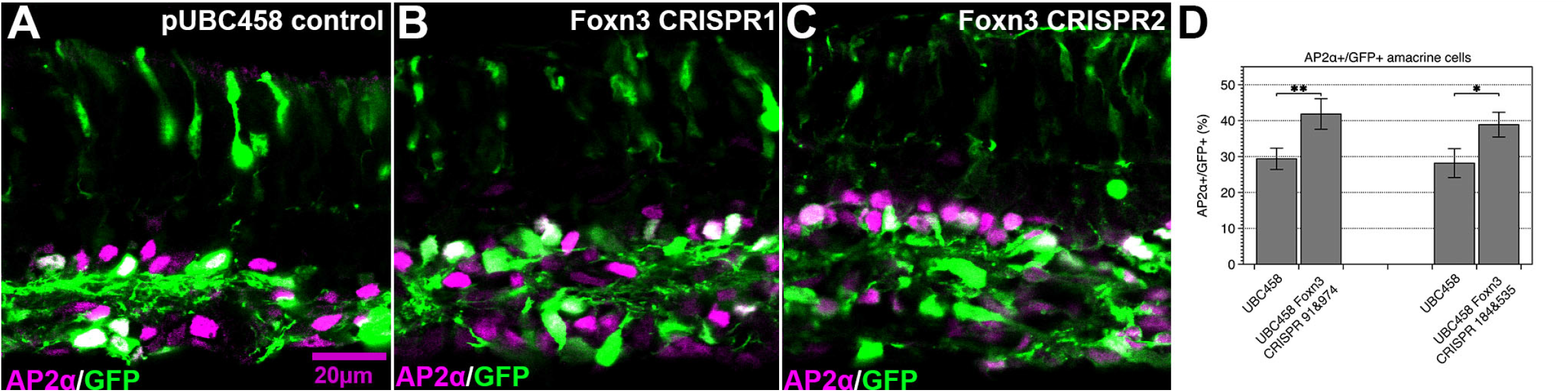
Disruption of Foxn3 by CRISPR/Cas9 increases amacrine cell formation in embryonic retinal explants. pUBC458 Cas9 or pUBC458 Cas9/Foxn3 sgRNA expression vectors, which express Cas9 and GFP, were introduced by electroporation into E16.5 retinas, which were then maintained in explant culture for 8 DIV. (A-D) Either of two Cas9 vectors with different sgRNAs targeting the Foxn3 gene increased the fraction of amacrine cells (AP2α, magenta) among GFP-labeled cells (green), relative to the control pUBC458 vector. Quantitation is shown in (D).* P < 0.05, ** P < 0.01. See Fig. 1 legend for labels.

To assess the consequences of Foxn3 overexpression in the developing retina, we introduced expression vectors for Foxn3 (without its 3’ UTR) and GFP into E16.5 retinas by electroporation, and then maintained the retinas in explant culture for 2 or 8 DIV. Forced expression of Foxn3 reduced the fraction of AP2α+ cells among GFP-labeled cells at 8 DIV, relative to the control expression vector (Fig. 7A,B,I). Ptf1a is transiently expressed in newly generated postmitotic amacrine cells (Fujitani et al., 2006). Two days after electroporation, the Foxn3 expression vector reduced the number of Ptf1a+ cells among the GFP-labeled cells, relative to the control (Fig. 7E-H, K). We also scored cell proliferation/DNA synthesis by incorporation of 5-ethynyl-2’-deoxyuridine (EdU) in the explants (Zeng et al., 2010). The fraction of EdU+ cells among the GFP-labeled cells was significantly increased at 2 DIV with the Foxn3 expression vector, relative to the control expression vector (Fig. 7C-D, J). These data indicate that Foxn3 can inhibit amacrine cell formation at an early step, prior to Ptf1a expression, and that expression of Foxn3 likely reduces cell cycle exit of retinal progenitor cells.

**Fig. 7.**
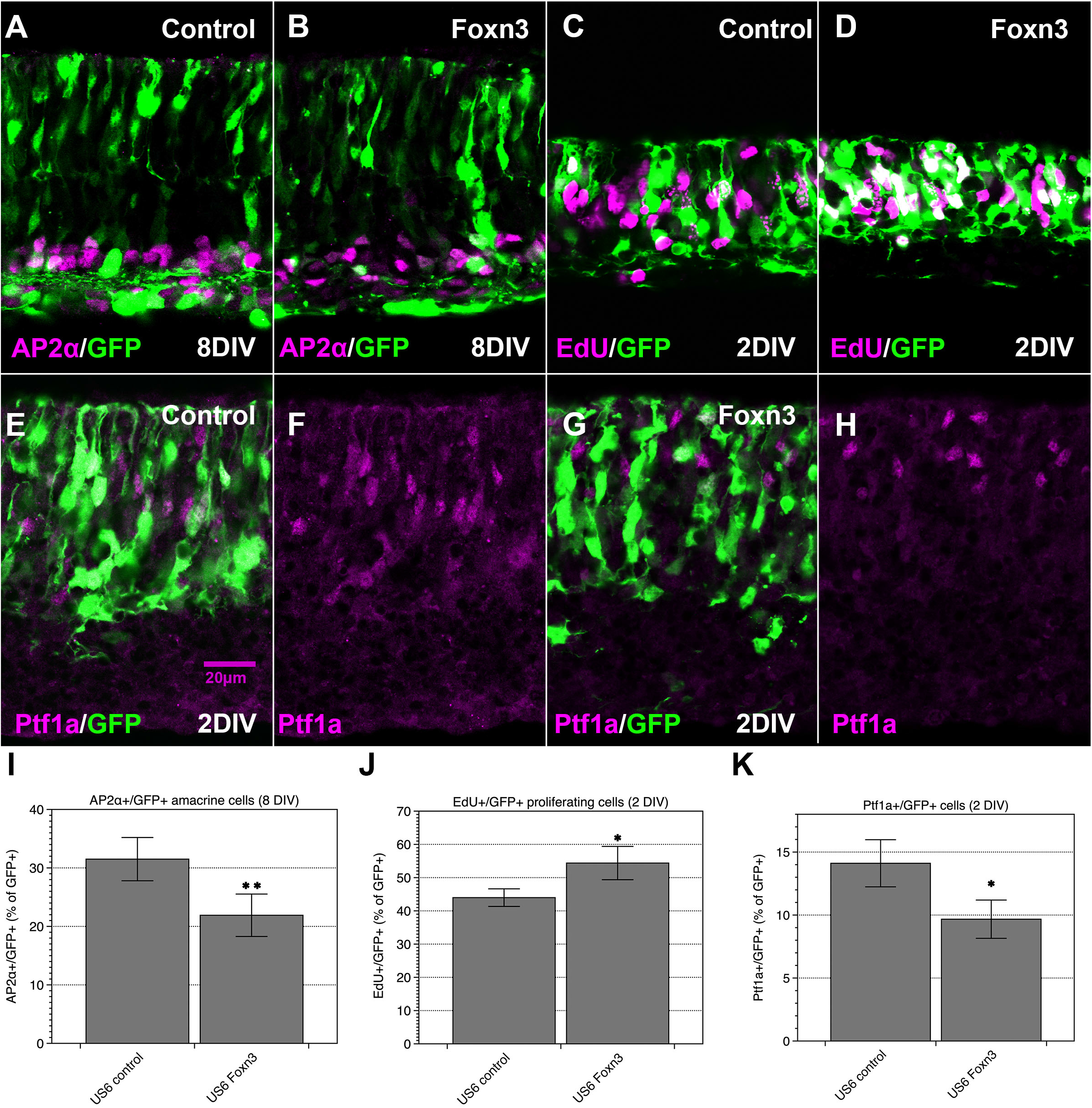
Ectopic expression of Foxn3 reduces amacrine cell formation and increases proliferation in the embryonic retinal explants. A GFP expression vector and either a Foxn3 expression vector or a control expression vector were introduced into E16.5 retinas by electroporation, which were then maintained in explant culture. (A,B) Expression of Foxn3 reduced the fraction of amacrine cells (AP2α, magenta) among GFP-labeled cells (green), relative to the control vector at 8DIV. (C, D) Foxn3 expression increased the fraction of GFP-labeled cells in S-phase, based on EdU incorporation (magenta) at 2DIV, relative to the control vector. (E-H) Foxn3 also reduced the fraction of GFP-labeled cells that express Ptf1 protein (magenta) at 2 DIV (G,H), relative to the control vector (E,F). (I-K) Quantitation: N=3 for each analysis. * P < 0.05, ** P < 0.01. Labels as in Fig. 1.

We tested whether co-expression of the Foxn3 protein could reverse the effects of forced miR-216b expression in the developing postnatal retina. We performed *in vivo* electroporation at P0 to co-deliver the miR-216b and Foxn3 expression vectors individually or together, as well as control vectors, into the retina. We assessed amacrine and bipolar cell markers in GFP-labeled cells at P12 (Fig. 8A-D, I, J). Expression of miR-216b increased the fraction of AP2α amacrine cells and decreased the fraction of Vsx2+ bipolar cells among GFP-labeled cells at P12, relative to the Luc shRNA control vector. Expression of Foxn3 reduced AP2α+ cells and increased Vsx2+ cells among the GFP-labeled cells at P12. Co-expression of Foxn3 with miR-216b partially reversed both the decrease in amacrine cells and the increase in bipolar cells among GFP-labeled cells that were observed with expression of miR-216b. We also observed an increase in GFP-labeled cells within the INL that were not labeled by either Vsx2 or AP2α when Foxn3 was expressed, with or without miR-216b (Fig. 8K). We also scored GFP-labeled cells in the ONL, which were likely to be rod photoreceptors based on morphology and lack of Vsx2 or AP2α expression (Fig. 8L). Expression of miR-216b did not significantly change the frequency of GFP-labeled cells in the ONL, although expression of Foxn3 slightly decreased the frequency of these cells, with or without co-expression of miR-216b. Finally, we scored Müller glial cells (Sox2 positive/AP2α negative) among the GFP-labeled cells in the INL (Fig. 8E-H, M). While expression of miR-216b did not significantly change the frequency of Müller glial cells among GFP-labeled cells, expression of Foxn3 modestly increased the frequency of Müller glial cells, and co-expression of miR-216b with Foxn3 reversed the increase. The ability of Foxn3 to partially reverse the effects of miR-216b are consistent with inhibition of Foxn3 as a component of the mechanism by which ectopic expression of miR-216b alters cell type generation in the developing retina.

**Fig. 8.**
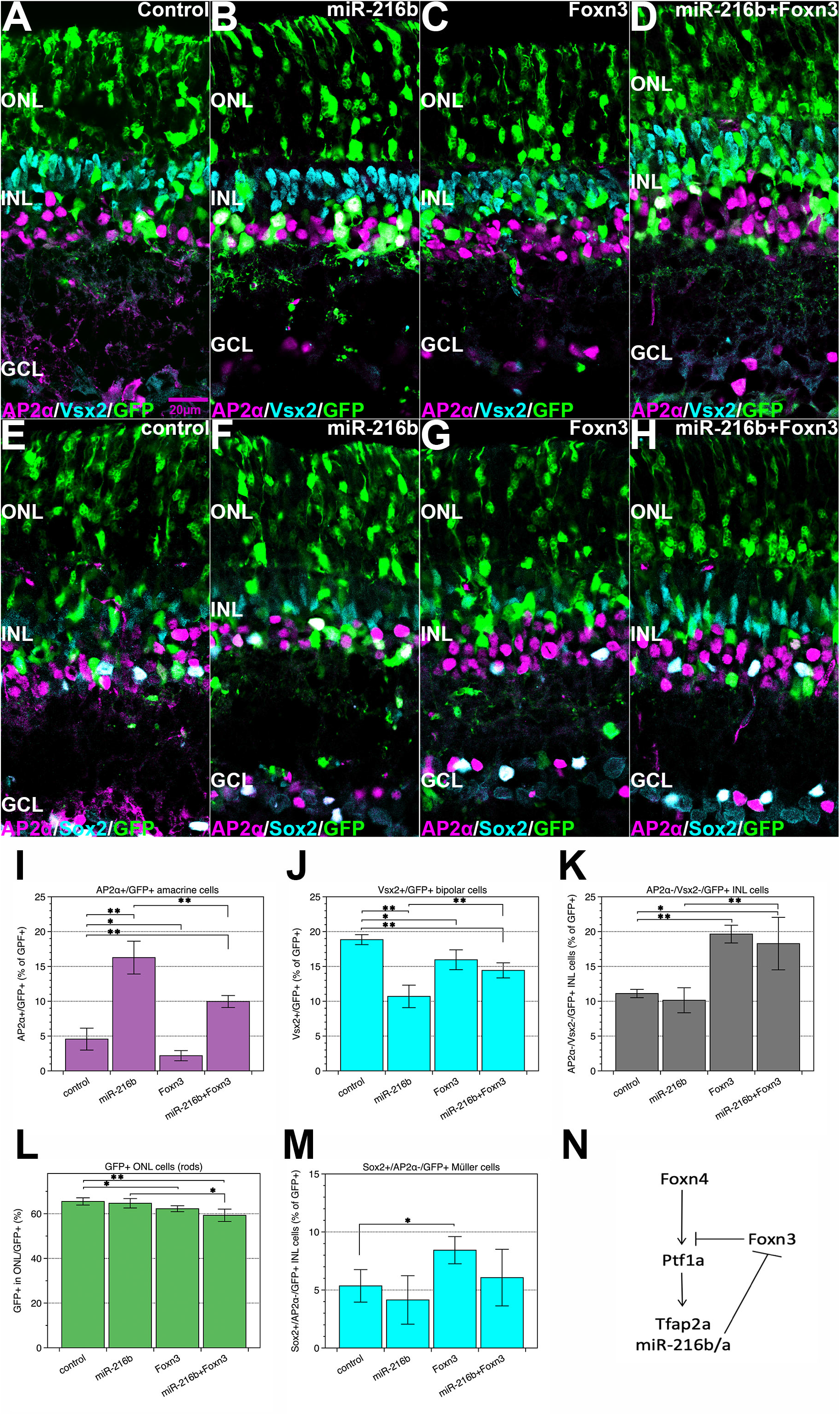
Co-expression of Foxn3 with miR-216b reduces the effects of miR-216b on amacrine cell and bipolar cell generation in the developing retina. A miR-216b and GFP vector or a control Luc shRNA and GFP vector, in combination with a Foxn3 or control expression vector, were introduced into retinas at P0 by *in vivo* electroporation. GFP-labeled (green) amacrine cells (AP2α, magenta) and bipolar cells (Vsx2, cyan) were assessed at P12 (A-D, J-L). The increased fraction of amacrine cells and decreased fraction of bipolar cells among GFP-labeled cells with miR-216b expression were partially reversed by the co-expression of Foxn3 (I, J). Foxn3 expression, with or without miR-216b, increased the fraction of GFP-labeled cells in the INL that were unlabeled by either AP2α or Vsx2 (K). The fraction of GFP-labeled cells in the ONL (presumptive rod photoreceptors) did not change with miR-216b expression, but was slightly reduced by Foxn3 expression, with or without miR-216b (L). (E-H, M) The fraction of GFP-labeled Müller glia (positive for Sox2, cyan; negative for AP2α, magenta) did not change with miR-216b expression, but was slightly increased by Foxn3 expression; the increase was reversed by co-expression of miR-216b (M). (N) Proposed model of miR-216b and Foxn3 function. (J-M) Quantitation: miR-216b vector N=4, others N=3. * P < 0.05, ** P < 0.01. Labels as in Fig. 1.

## Discussion

Expression of miR-216b-5p overlaps with differentiated amacrine cells in the mouse retina. miR-216b is part of a genomic miRNA cluster with miR-216a and miR-217, two miRNAs expressed from a shared transcription unit (Kato et al., 2009). All three miRNAs are expressed in the mouse inner retina, consistent with previous observations of miR-216a/miR-217 expression in developing zebrafish retinas (Olena et al., 2015; Wienholds et al., 2005), and these miRNAs were downregulated in retinas in the absence of the Ptf1a, a bHLH transcription factor required for the differentiation of nearly all horizontal and amacrine cells. Since Ptf1a is transiently upregulated only after amacrine cells become postmitotic (Fujitani et al., 2006), all three miRNAs are preferentially expressed in postmitotic amacrine cells. The lack of expression of these miRNAs in the neuroblast layer of the developing retina, as well as their dependence on Ptf1a for high level expression, suggests that miR-216a/b are unlikely to regulate the initial commitment or cell cycle exit of amacrine cells during retinal development. Nonetheless, ectopic expression of pre-miR-216a or pre-miR-216b in the developing mouse retina led to a significant increase in amacrine cells. The increase in amacrine cells with miR-216b depended on the miR-216b-5p seed sequence, indicating that seed-based target repression by mature miR-216b-5p is likely to drive the increase in amacrine cells. However, expression of miR-216b in retinas in which the Ptf1a gene was disrupted failed to generate amacrine cells, indicating that expression of miR-216b cannot replace Ptf1a. We propose that the normal role of miR-216b and miR-216a is likely to be reducing the expression of repressors of amacrine cell differentiation (and possibly other genes) in postmitotic, differentiating amacrine cells, and that ectopic expression of these miRNAs in retinal progenitor cells leads to premature inhibition of their targets and a consequent increase in amacrine cell formation. A potential functional requirement for the miR-216a/miR-216b/miR-217 genomic cluster in the retinal development has not been evaluated, as the homozygous deletion of the gene cluster in mouse embryos is lethal prior to retina formation (Azevedo-Pouly et al., 2017). However, mice lacking the individual miRNAs are viable, suggesting that miR-216a/b may have redundant roles in embryonic development (Sutaria et al., 2019). In zebrafish, miR-216a has been linked to regulation of the Notch pathway by sorting nexin 5 (SNX5) during development (Olena et al., 2015), but the miR-216a target sites are not conserved in mouse Snx5. Zebrafish miR-216a also is involved in Müller cell reprogramming during retinal regeneration (Kara et al., 2019).

We identified Foxn3 as a target of miR-216a/b-5p, based on crosslinking of the Argonaute protein by PAR-CLIP near an evolutionarily conserved miR-216a/b-5p seed match in the Foxn3 3’ UTR, and we confirmed that miR-216b-5p could repress a reporter containing a region from the Foxn3 3’ UTR via two miR-216a/b-5p seed matches. Strikingly, inhibition of Foxn3 expression in the postnatal retina by RNAi, or disruption of the Foxn3 gene by CRISPR/Cas9 in embryonic retinal explants, led to an increase in amacrine cell formation, similar to that observed with miR-216b overexpression. Furthermore, ectopic expression of Foxn3 in retinal explants reduced the number of differentiated amacrine cells. Ectopic Foxn3 also reduced expression of Ptf1a and modestly increased the fraction of proliferating cells when assayed two days after electroporation, consistent with inhibition of amacrine cell formation at an early step, prior to cell cycle exit. Taken together, these data indicate that Foxn3 is a negative regulator of amacrine cell formation in the developing retina, likely limiting the initial formation of amacrine cells. Co-expression of Foxn3 with miR-216b partially reversed the increase in amacrine cell numbers seen with miR-216b expression, suggesting that inhibition of Foxn3 likely contributes to the effects of ectopic miR-216b. However, additional miR-216a/b targets also may contribute to the phenotypes observed after ectopic expression of miR-216a/b. In addition, regulation of Foxn3 by other miRNAs has been described in non-neural cells, and a previously validated target site for miR-135b-5p is located near the miR-216a/b site in the 3’ UTR of the Foxn3 mRNA (Han et al., 2019). miR-135a-5p and miR-135b-5p are expressed in the developing retina (Supplemental Table 1), so additive or redundant regulation of the Foxn3 mRNA by multiple miRNAs in the retina is possible.

Foxn3 has been linked to multiple biological processes, including cell proliferation, epithelial-mesenchymal transition, and regulation of glucose metabolism (Huot et al., 2014; Karanth et al., 2018; Li et al., 2017). Reduced Foxn3 function in frogs or mice leads to defects in head neural crest and a small eye phenotype (Samaan et al., 2010; Schuff et al., 2007). We focused on Foxn3 both because the conserved miR-216a/b-5p seed match in the Foxn3 3’ UTR had the most crosslinks among potential miR-216a/b-5p sites in the retina Argonaute PAR-CLIP data, and because Foxn3 is closely related to Foxn4. Foxn4 is a transcription factor expressed in proliferating retinal progenitor cells that is required for amacrine and horizontal cell formation (Li et al., 2004), and this role is evolutionarily conserved (Boije et al., 2013). Foxn4 is a transcriptional activator that directly activates transcription of Ptf1a and other genes (Del Barrio et al., 2007; Liu et al., 2020; Luo et al., 2012). Foxn3 and Foxn4 have been reported to bind to similar DNA target sequences, although Foxn3 also recognizes an additional class of DNA target sequences that Foxn4 does not (Nakagawa et al., 2013; Rogers et al., 2019). Since Foxn3 is a transcriptional repressor (Scott and Plon, 2005), we speculate that Foxn3 could bind to and repress direct Foxn4 target genes, thus reducing amacrine cell formation. Consistent with this model, overexpression of Foxn3 reduced the number of Ptf1a positive cells in the developing retina. Alternately, the ability of Foxn3 to recognize binding sites structurally distinct from those sites recognized by Foxn4 could allow Foxn3 to regulate other target genes.

Ectopic expression of miR-216b or inhibition of Foxn3 by RNAi significantly reduced the number of retinal bipolar cells among the electroporated retinal cells, while co-expressed Foxn3 partially reversed the reduction of bipolar cells by miR-216b. While clones of mouse retinal cells derived from a single progenitor cell can contain both amacrine and bipolar cells (Turner et al., 1990), the majority of these two cell types are generated at different times during development, with the peak of amacrine formation near E17, and the peak of bipolar generation at P1-P3 (Voinescu et al., 2009). Ectopic expression of Foxn4 during retinal development reduces bipolar cell formation (Li et al., 2004). Lineage tracing of retinal cells in Dll4-Cre mice indicates that most or all bipolar cells are generated from retinal progenitor cells that did not express Dll4. The Dll4 gene is directly activated by Foxn4 (Luo et al., 2012; Zou et al., 2015), suggesting that bipolar cells arise from progenitor cells in which Foxn4 target genes were not activated. These observations suggest that both increased amacrine cell formation and decreased bipolar cell formation could arise from elevated Foxn4 activity, consistent with a model in which Foxn3 limits Foxn4 function, and miR-216b-5p inhibits Foxn3 expression (Fig. 8N). We also observed that Foxn3 mRNA is expressed in differentiated bipolar cells in the P12 retina, consistent with a role for Foxn3 in bipolar cells. However, overexpression of Foxn3 by itself in postnatal retinas reduced the number of bipolar cells relative to control, albeit less than miR-216b, suggesting that Foxn3 can have other effects, directly or indirectly, on bipolar cell formation. It remains possible that other targets of miR-216b-5p could contribute to the reduced number of bipolar cells. We also observed that ectopic Foxn3 expression could modestly increase Müller glial cell frequency and slightly decrease the frequency of presumptive rods, while ectopic miR-216b expression did not. Co-expression of miR-216b with Foxn3 prevented the increase in Müller glial cells, suggesting that additional targets of miR-216b interact with Foxn3.

Forced expression of Foxn3 in retinal progenitor cells did not prevent all amacrine cell formation among the electroporated cells. Foxn3 is known to interact with several other proteins (Li et al., 2017; Scott and Plon, 2005), so Foxn3 function may be limited by the absence of a cofactor in some retinal cells. Function or stability of the Foxn3 protein also could be modulated by post-translational modification, as has been reported for the Foxn2 protein, a closely related transcriptional repressor (Ma et al., 2018). However, we cannot rule out that the Foxn3 vector was expressed at an insufficient level in some cells, or that some electroporated retinal cells may have committed to the amacrine cell fate prior to ectopic Foxn3 expression.

Morphological and molecular analyses have identified numerous subtypes of amacrine cells in the mouse retina (MacNeil and Masland, 1998; Macosko et al., 2015; Yan et al., 2020). Based on recent single cell sequencing analysis of amacrine cells (Yan et al., 2020), cholinergic amacrine cells express Foxn3 mRNA. Since cholinergic amacrine cells are generated prior to E17 (Voinescu et al., 2009), the RNAi and CRISPR experiments were too late to reduce Foxn3 expression in these cells. It is also possible that miR-216a/b limit the expression of the Foxn3 protein in cholinergic amacrine cells.

Previous analyses of miRNA function in the retina have revealed roles in developmental timing. Loss of all miRNAs after disruption of the Dicer processing enzyme can lead to prolonged production of early retinal cell types and loss of later cell types, including amacrine cells (Georgi and Reh, 2010). Our observation that ectopic expression of miR-216b can lead to the generation of additional amacrine cells and loss of bipolar cells is consistent with a role for miRNAs in developmental timing of the retina. Other miRNAs also have been implicated in amacrine cell development. We previously identified miR-181b as another miRNA with elevated expression in mouse amacrine cells that depends on Ptf1a function (Zhuang et al., 2020). In Medaka, the levels of miR-181a/b expression in retina depends on TGF-beta signaling, and miR-181a/b have been implicated in amacrine and ganglion cell process maturation (Carrella et al., 2015a; Carrella et al., 2015b).

Transcription of the miR-216a/217 miRNA primary transcript in diabetic kidney cells has been reported to be activated by TGF-beta signaling and bHLH transcription factors (Kato et al., 2009). Intriguingly, amacrine cells express TGF-beta2, as well as the TGFBR2 receptor (Ma et al., 2007; Tachibana et al., 2016) which could potentially contribute to miR-216a/b/miR-217 upregulation via autocrine signaling. In addition to Ptf1a, the Ascl1, Neurod1, and Neurod4 bHLH proteins are redundantly required for amacrine and horizontal cell formation in retina (Akagi et al., 2004; Inoue et al., 2002), suggesting that those proteins could regulate transcription of miR-216b/a/miR-217 in the retina. A genome wide analysis of Ptf1a and Ascl1 binding sites identified ChIP-Seq peaks for Ptf1a and Ascl1, including in the region upstream of the miR-216b miRNA precursor on Chr.11 in cells from the mouse spinal cord (Borromeo et al., 2014). In addition, miR-216a/b and miR-217 are highly expressed in mouse pancreas (Azevedo-Pouly et al., 2017; Sutaria et al., 2019), a tissue dependent on Ptf1a function during development, indicating a link between Ptf1a function and expression of these miRNA genes in multiple tissues.

Our results identify Foxn3 as a novel regulator of amacrine cell and bipolar formation during mouse retinal development, and they suggest a role for miR-216b in gene regulation in amacrine cells. Both of these factors are expected to function as repressors of their targets, suggesting that their roles are likely to be to limit the expression of other factors. Multiple repressive factors, such as Prdm1 and Vsx2, have been implicated in the regulation of bipolar cell and amacrine cell formation (Goodson et al., 2020), as well as additional, unidentified transcriptional repressors in bipolar or Müller cells (Chan et al., 2020). Future experiments that determine the targets of Foxn3 in the retina should help to decipher the regulatory networks that control cell diversification in the developing retina.

## Materials and Methods

### Plasmid construction

Plasmids were constructed using standard techniques. Oligonucleotides used for plasmid construction are show in Supplemental Table S6. The pUS2, pUS2-MT, pUS2-eGFP, pUS2-Puro, pUS2-Luc, pUI4-GFP-SIBR, pUI4 Luc-shRNA (Chung et al., 2006) and pUS2-Tol2 transposase (Gupta et al., 2018) have been described previously. The pUS6, pUI6-GFP-SIBR, and pUI6 Luc-shRNA plasmids are variants of pUS2, pUI4-GFP-SIBR, and pUI4 Luc-shRNA that have ITRs from the Tol2 transposon flanking the expression cassettes to allow for integration in the presence of Tol2 transposase (Balciunas et al., 2006). Partial primary transcript sequences (pre-miRNAs and some flanking sequences) for the mouse miR-216a, miR-216b, and miR-217 genes were amplified by PCR from mouse genomic DNA and cloned into pUI4-GFP-SIBR and pUI6-GFP-SIBR using primers with the indicated restriction sites. Pre-miR-216b-mut, with a 3-nt mutation in the seed sequence of miR-216b-5p (AATCTCT to AATAGAT) and a compensating change in miR-216b-3p to maintain base pairing in the pre-miRNA, was synthesized as a gBlock (IDT) and inserted into pUI6-GFP-SIBR. The mouse Foxn3 and Foxn4 coding regions were isolated by RT-PCR from embryonic mouse retina RNA and inserted into pUS2 and pUS6. Foxn3 shRNA constructs pUI6-GFP-SIBR Foxn3-195×2 and pUI6-GFP-SIBR Foxn3-635×2, contain two tandem miR-155-based SIBR cassettes (Chung et al., 2006), which target the coding region of the mouse Foxn3 mRNA. NanoLuc was PCR amplified from plasmid pNL.1.1 (Promega) and cloned into pUS2-MT to generate pUS2-MT-Nanoluc. Partial Foxn3 3’UTR fragments containing wild type or mutated miR-216a/b target sites (sequence altered to match the seed sequence of miR-216b-mutant) were amplified by RT-PCR from mouse retina RNA, and were inserted downstream of firefly Luciferase in pUS2-Luc. pUBC458op was made by replacing CBH promoter in pX458 (AddGene #48138)(Ran et al., 2013) with the UBC promoter and first intron from pUI4-GFP-SIBR, followed by replacement of the original sgRNA scaffold with an optimized sgRNA scaffold (Dang et al., 2015), retaining the U6 promoter and Bbs1 cloning sites from pX458. Oligonucleotides for Foxn3 sgRNAs were designed using E-CRISP (Heigwer et al., 2014) and inserted into pUBC458op. A second U6 sgRNA expression cassette was inserted adjacent to the existing U6-sgRNA cassette to express two sgRNAs.

### Mice and processing of retinas

All animal experiments were approved by the Institutional Animal Care & Use Committee at the University of Michigan. Wild-type retinas were isolated from CD-1 mice (Charles River). Ptf1a mutant mice have the Ptf1a coding region replaced with the Cre coding region (Kawaguchi et al., 2002). Eyes were fixed with 2% paraformaldehyde in phosphate buffered saline (PBS) for 30 min at room temperature, retinas were dissected from the eyes, then cryoprotected in PBS containing 30% sucrose for 2 hours at 4°C, and embedded in OCT, prior to preparation of cryosections (16 μm) using a cryostat. All the retinas were processed similarly to obtain tissue for in situ hybridization or antibody staining.

### In vivo and in vitro electroporation, and retina explant culture

1 μl of plasmid DNA vectors were injected into the subretinal space of P0 mouse pups and introduced into cells by *in vivo* electroporation as described (Zhang et al., 2012). The concentration and combination of DNA vectors are as listed below. 2 μg/μl pUI4 miRNA expression vectors or pUI4 Luc-shRNA control were injected in Fig.2. 2 μg/μl pUI6 miR-216b expression vector, 2 μg/μl pUI6 miR-216b-mut, 2 μg/μl pUI6 Luc-shRNA, or 2 μg/μl pUI6 Foxn3 RNAi vectors, and 200 ng/μL pUS2-Tol2 were injected in Fig.5. 2 μg/μl pUI6 miR-216b expression vector or 2 μg/μl pUI6 Luc-shRNA, and 1 μg/μl pUS6-Foxn3 expression vector or 1 μg/μl pUS6 control, and 300 ng/μL pUS2-Tol2 were injected in Fig.8. After DNA injection, five square 80-V pulses of 50-ms duration with 950-ms intervals were applied by using an ECM830 (BTX) with forcep-type electrodes. Electroporated retinas were collected at P7 (Fig. 2) or P12 (Fig.5 and Fig.8).

For retinal explants, heads from E16 mouse embryos were split in half along the sagittal suture and the cut surface of the tissue was placed on a square plate electrode. Plasmids were introduced into retinal cells by *in vitro* electroporation. 0.5 μl of plasmid DNA vectors were injected into the subretinal space of each eye. 2 μg/μl of pUBC458op Foxn3 CRISPR vectors or pUBC458op control (Fig.6), 2 μg/μl pUI6 miR-216b or pUI6 Luc shRNA expression vectors and 200 ng/ μl pUS2-Tol2 (Supplemental Fig.S3), or 1 μg/μl pUS6 or pUS6-Foxn3 overexpression vector combined with 1 μg/μl pUS6-GFP expression vector and 200 ng/μL pUS2-Tol2 (Fig.7) were injected. After DNA injection, five square 30-V pulses of 50-ms duration with 950-ms intervals were applied by using an ECM830 (BTX) with a platinum needle electrode above the eye in combination with the plate electrode underneath the embryonic tissue (Matsuda and Cepko, 2004). Electroporated retinas were dissected from the eyes and placed on Transwells with polyester membrane (Corning, pore size 0.4 μm), with the ganglion cell layer facing upwards. The Transwells were inserted into six-well plates and cultured in 1.2 ml of high-glucose Dulbecco’s modified Eagle’s medium (DMEM, Thermofisher) supplemented with 20% horse serum and 1% penicillin-streptomycin (Thermofisher). Retinas were collected at 2 or 8 days after electroporation.

### In situ hybridization and Immunofluorescence

Detection of miRNAs by in situ hybridization with RNA probes was as described previously (Deo et al., 2006; Zhuang et al., 2020), using gel purified oligoribonucleotide probes synthesized with both 5′ and 3′ fluorescein modifications (Sigma). Fluorescein labeled probes were detected using an alkaline-phosphatase conjugated anti-fluorescein Fab2 antibody (Roche), followed by BCIP/NBT (Roche) staining, or Fast Red (Sigma) staining. For Fast Red staining, after alkaline-phosphatase conjugated anti-fluorescein Fab2 antibody staining, slides were washed in 1XTBS for 10 min twice. The Fast Red tablet was reconstituted in water according to the manufacture’s instruction and 200 μl fast red solution was added to each slide. Slides were monitored closely, and staining was stopped by washing the slides in PBS. Sequences of oligoribonucleotide probes are in Supplemental Table S6.

mRNA *in situ* hybridization with RNA probes was performed as described previously (Deo et al., 2006; Zhang et al., 2012). Restriction enzyme linearized plasmid pUS2-Foxn3 or pUS2-Foxn4 was used as template for the synthesis of UTP-fluorescein or UTP-digoxigenin labeled riboprobes by *in vitro* transcription. mRNA of Foxn3 or Foxn4 was detected with split initiator mRNA *in situ* HCR (Choi et al., 2018). Each mRNA was hybridized with a pool of 20 probe pairs (IDT), and followed by using matching 72 nt hairpin amplifiers (Molecular Technologies). Detailed protocols for mRNA *in situ* HCR were described previously (Zhuang et al., 2020). Sequences of *in situ* HCR probes are in Supplemental Table S6.

Indirect immunofluorescence analysis was performed on retinas without *in situ* hybridization or after *in situ* hybridization. For AP2α antibody staining, retina sections were blocked with 1:400 unlabeled affinity purified Fab fragment donkey anti-mouse IgG (H+L) and 5% donkey serum in PBS with 0.05% TritonX-100 at 4°C overnight, incubated with mouse anti-AP2α (1:1000, Developmental Studies Hybridoma Bank), then incubated with Alexa Fluor 647-conjugated or Alexa Fluor 546-conjugated anti-mouse secondary antibody (both 1:1000, Jackson Immunoresearch) for one hour at room temperature. For Vsx2 (Chx10), Ptf1a and Sox2 staining, tissue sections were blocked with 5% donkey serum, and incubated with sheep anti-Chx10 (1:100, Millipore), rabbit anti-Ptf1a (1:800, Li and Edlund, 2001), or rabbit anti-Sox2 (1:1000, Millipore) at 4°C overnight, followed by incubation with Alexa 594-conjugated anti-sheep secondary antibody (1:1000, Jackson Immunoresearch), or Alexa 594-conjugated anti-rabbit secondary antibody (1:1000, Jackson Immunoresearch). After antibody staining, Hoechst 33342 (Roche) was used to stain the nuclei. For EdU labeling, retinal explants were incubated with EdU for 12 hrs prior to collection, then the Click-iT EdU Cell Proliferation kit (ThermoFisher) was used to detect EdU, as directed by the manufacturer.

### Microscopy and imaging

BCIP/NBT staining was imaged with a Leica DSMIRB microscope with a SPOT-RT digital camera, or a Zeiss Axiovert microscope with an AmScope MU200 camera. Fluorescent antibody staining with GFP or in situ hybridization/HCR were imaged using an Olympus FV1000 confocal microscope. Images were analyzed and colors were assigned with FV10-ASW 3.1 Viewer or FIJI/ImageJ. Representative images were cropped, assembled, and labeled using Adobe Photoshop.

### Quantitation, graphs, and statistics

GFP+ /antibody positive cells were analyzed from at least 3 retinas for each condition. Cells were counted manually from images, and analysis was not blinded. Cell counts were totaled across several sections for each sample. Error bars show standard deviation for all graphs. T-tests: 1 tail, two samples with equal variance. Graphs were generated using DataGraph (Visual Data Tools, Inc.) or Microsoft Excel.

### qRT-PCR

PCR primers are listed in Supplemental Table S6. RNA was extracted from retina using TRIzol (ThermoFisher) and treated with RNase-free DNaseI (Promega). For miRNA qRT-PCR, 1 μg of RNA was polyadenylated using a Poly (A) Tailing Kit (Thermofisher). After re-purification with TRIzol reagent, total RNA was reverse-transcribed with 0.5ug of poly (T) adaptor containing a 5’-end universal tag sequence and SuperScript™ II Reverse Transcriptase (Thermofisher). Specific miRNA primers, in combination with a universal reverse primer that binds to the universal tag sequence, were used to amplify specific miRNA sequences (Thomson et al., 2006). For mRNA qRT-PCR, 1 μg of RNA was reverse transcribed using Superscript II and random 12mer primers. 2.5 ng of cDNA for miRNA assay, or 20 ng of cDNA for mRNA assay was used per real-time PCR with SYBR Green (Bio-Rad) to detect DNA, using a BioRad iCycler with MyiQ light module. Expression of Foxn3 mRNA was normalized to the GAPDH mRNA. Data presented are mean of three RNA samples for each condition, each assayed using triplicate PCR replicates. The 2^−ΔΔ**CT**^ method was used to calculate fold change.

### Transfections, Luciferase assays, and cell culture RNA collection

HEK293 cells were grown in DMEM with 10% fetal bovine serum (FBS) and 1:100 penicillin-streptomycin (ThermoFisher). In 24-well plates, 50 ng pUS2-Luc, pUS2-Luc-Foxn3-UTR-WT, pUS2-Luc-Foxn3-UTR-3’-216b-mutant, pUS2-Luc-Foxn3-UTR-5’-216b-mutant, or pUS2-Luc-Foxn3-UTR-5’+3’-216b-mutant were cotransfected with 250ng of UI6, UI6-miR-216b, or pUI6-miR-216b-mutant into each well. 25ng of pUS2-MT-Nanoluc was also contransfected into each well as transfection control. Transfections were performed with Lipofectamine 2000 (ThermoFisher) according to the manufacturer’s instructions. Reporter activity was assayed 48 hr after transfection using the Nano-Glo Dual Luciferase Assay kit (Promega). Firefly Luciferase activity was normalized to Nanoluciferase activity to control for transfection efficiency variation. A FLUOstar OPTIMA plate reader was used to measure luminescence intensity. Data presented are averaged from three independent transfections for each condition, assayed using duplicate luciferase assay replicates.

Mouse P19 cells were grown in MEM-α (ThermoFisher) with 2.5% FBS, 7.5% calf serum and 1:100 penicillin-streptomycin. To determine the effectiveness of Foxn3 RNAi constructs in P19 cells, 3μg of Foxn3 RNAi vectors and 1μg of pUS2-Puro were cotransfected into one well of a 6-well plate. Transfections were performed with TransIT-LT1 Transfection Reagent (Mirus) according to the manufacturer’s instructions. The transfected cells were shifted to medium with 12.5 μg/ml puromycin 8 hours after transfection and selected for 16 hours. Total RNA was extracted by using TRIzol 48 hours after transfection.

### PAR-CLIP Libraries

Argonaute PAR-CLIP libraries were prepared based on the method described by Hafner et al. (Hafner et al., 2010), with modifications based in part on tissue processing used for HITS-CLIP (Chi et al., 2009). P0 retinas from CD-1 mice were isolated and grown as explants for 24 hours in high glucose DMEM supplemented with 20% horse serum and 0.1 mM 4-thiouridine (Sigma). Six retinas were used to make one library. Live, intact retina explants were irradiated with 365 nm UV light (~300mJ/cm2) using a Stratalinker (Stratagene), then lysed in 300 μl of low salt lysis buffer (50 mM HEPES pH 7.5, 50 mM KCl, 0.5% sodium deoxycholate, 0.1% SDS, 0.5% NP-40, 0.5 mM DTT and Protease inhibitor cocktail (Sigma)) with 6 μl of RNaseout (Life Technologies). The lysates were sequentially treated with 12.6 μl of RNase-free DNase (Promega) and 7.5 μl of 1:1330 diluted micrococcal nuclease (11.3 gel units, New England Biolabs: NEB) for 5 min at 37 °C. 8.4 μl of 0.5 M EGTA (Sigma) was added to stop the micrococcal nuclease digestion. Lysates were cleared by centrifugation at 14000 rpm for 10 min at 4°C. Cleared lysates were incubated with 10 μg of 2A8 antibody (EMD-Millipore) (Nelson et al., 2007) bound to Protein G Dynabeads (Life Technologies) and rotated at 4°C overnight. Beads were washed 3 times in IP wash buffer (50 mM HEPES pH 7.5, 300 mM KCl, 0.5% NP-40, 0.5 mM DTT and Protease inhibitor cocktail), 3 times in high salt wash buffer (50 mM HEPES pH 7.5, 500mM KCl, 0.5% NP-40, 0.5 mM DTT and Protease inhibitor cocktail), and resuspended in dephosphorylation buffer (100 mM NaCl, 50 mM Tris-HCl pH7.9, and 10 mM MgCl_2_). 4 μl of Calf intestinal alkaline phosphatase (NEB) was added to dephosphorylate the RNA for 15 min at 37 °C. Beads were washed twice in phosphatase wash buffer (50 mM Tris-HCl pH7.5, 20 mM EGTA, 0.5% NP-40), and twice in polynucleotide kinase (PNK) buffer (50 mM Tris-HCl pH7.5, 50 Mm NaCl, 10 mM MgCl2). Beads were resuspended in 80 μl of PNK buffer with 5mM DTT, and incubated with 8 μl of T4 Polynucleotide Kinase (NEB) and 2 μl of 10 mM ATP for 30 min at 37 °C. The beads were washed 5 times in PNK buffer. The protein-RNA complexes were eluted in 30 μl of 1X Novex Tris-glycine SDS sample buffer (Life Technologies), and separated by 10% SDS-PAGE. A gel slice corresponding to 100 to 150 kd was isolated (based on size markers in an adjacent lane; Precision Plus Protein™ Dual Color Standards from Bio-Rad). The gel slice was transferred to a D-Tube Dialyzer Midi Tube (EMD-Millipore). The protein-RNA complexes were electroeluted in 1x NuPage MOPS SDS running buffer (Life Technologies) at 100 V for hr. The eluate was digested in 50 mM Tris-HCl pH7.5, 72 mM NaCl, 6 mM EDTA, 1 mM SDS, and 1.2 mg/ml Proteinase K (Sigma) at 55 °C for 30 min. RNA was recovered by acidic phenol/chloroform extraction and ethanol precipitation. The recovered RNA was used to prepare a cDNA library using a TruSeq Small RNA Library Preparation Kit (Illumina) according the manufacturer’s directions, with the following modification: after reverse transcription, the reaction was denatured, separated on a 10% TBE urea gel; a gel slice corresponding to 80 to 150 nt was isolated (based on size markers in an adjacent lane: low range ssRNA ladder, NEB). The reverse transcribed product in the gel slice was eluted, precipitated by adding 2-propanol, suspended in Tris-HCl buffer, then used in subsequent steps. PAR-CLIP libraries were sequenced on an Illumina HiSeq 2500 (University of Michigan Advanced Genomics Core).

Single-end read PAR-CLIP cDNA sequences were mapped to the mouse mm10 genome using STAR (Dobin and Gingeras, 2016). Sequences mapping to ChrM or rRNA sequences were removed. Multiple reads with identical sequence and length were considered duplicates and only a single read was retained. We considered sequence as well as length to avoid discarding potentially identical cDNA fragments with crosslinks at different positions, although this meant that duplicate reads with distinct errors introduced during the PCR amplification of the library would be considered as separate reads. T to C sequence substitutions relative to the genome were identified with Bambino (Edmonson et al., 2011). Reads with low base quality mismatches or near the ends of reads were filtered, so the count of T to C substitutions and coverage at a specific position is conservative (both crosslink counts and read coverage at crosslink sites exclude some input sequences due to filters). T to C substitutions present at the same genomic position in at least two of the five Argonaute PAR-CLIP libraries, but not present in the two IgG control libraries, were identified, combined, and filtered using scripts, BEDtools (Quinlan, 2014), and an in-house program. Deletions at the same position as consistent T to C substitutions were also considered crosslinked reads. Consistent crosslinks within 15nt of a position with a greater number of reads with a consistent crosslink were removed, to create the set of predominant crosslinks. Predominant crosslink sites were annotated using GENCODE VM16 gene/exon annotations (Frankish et al., 2019) and BEDTools. 6 or 7nt miRNA seed matches with the 5’ base of the match within +/− 15nt of a predominant crosslink site (31nt windows) were identified, based on seeds from miRNAs detected in P0 retinas (Supplemental Table S2). To identify evolutionary conservation of miRNA seed sites, PhastCons (Siepel et al., 2005) scores were determined for both the 31nt window and individual seed matches at each predominant crosslink, based on the mm10 MAF60 alignment. In addition, sequences corresponding to the seed sequences from several selected species were retrieved via hg19 MAF100 multiple alignments, using Galaxy (Afgan et al., 2018) at the public server usegalaxy.org. The subset of predominant crosslinks that had a 6 or 7nt match to the miR-216a and/or miR-216b seed sequences in the genome were selected for further analysis. Plots of read coverage were generated using IGV (Thorvaldsdottir et al., 2013).

To evaluate miRNA seed enrichment near predominant crosslinks (Supplemental Fig. 4), 3’ UTR exons with crosslinks were identified. Overlapping exons on the same strand were merged, and crosslinks near the ends of exons were filtered to avoid seed matches outside of the exons. All seed matches within +/− 15 nt were counted. If overlapping matches of different lengths (e.g. 6 nt vs 7 nt) were possible for the same seed match, only the longest match was counted. To determine the expected distribution of miRNA seeds near crosslink positions, a Monte-Carlo method was used. Randomly selected T’s from the same set of exons were used as pseudo-crosslinks (same number of pseudo-crosslinks selected per exon as predominant crosslinks identified for that exon, with same filters on position). Seed matches were counted within +/− 15nt of the pseudo-crosslinks, for 100,00 trials. Bonferonni correction was used to adjust P values for multiple testing.

### Small RNA libraries

Retinas from individual E18.5 embryos (Ptf1a knockout or heterozygous) were isolated and pooled. The genotype of embryos was determined by PCR using the primers: Ptf1a-F CGAGGACAACGTCAGCTATTG, Ptf1a-R TCTCGCACATTAGCGGCTTG, Cre-F CATGCTTCATCGTCGGTCC, and Cre-R GATCATCAGCTACACCAGAG. Total RNA was purified by using TRIzol, then denatured and separated on a 15% TBE urea gel. Small RNA was recovered from a gel slice corresponding to approximately 18 to 30 nt (based on size markers in an adjacent lane: 20/100 Ladder, IDT). The gel purified small RNA from 1 μg of total RNA was used to prepare a small RNA library using a TruSeq Small RNA Library Preparation Kit (Illumina) according the manufacturer’s directions, with the following modification: after reverse transcription, the reaction was denatured, separated on a 10% TBE urea gel; a gel slice corresponding to 60 to 80 nt was isolated (based on size markers in an adjacent lane: low range ssRNA ladder, NEB). The reverse transcribed product in the gel slice was eluted, precipitated by adding 2-propanol, suspended in Tris-Hcl buffer, and then used in subsequent steps. Small RNA libraries were sequenced on an Illumina HiSeq 2500 (University of Michigan Advanced Genomics Core). Insert sequences had adaptor sequences trimmed and inserts <18 nt or >30nt were removed. Sequences were then mapped to mature miRNA sequences in miRBase 22 (www.mirbase.org)(Kozomara et al., 2019). Ambiguous mappings and unmapped sequences were discarded. Read counts were normalized to input library size. We also determined the most frequent sequence for each miRNA. Small RNA libraries from P0 CD-1 mouse retinas were made and analyzed by a similar protocol, except that the NEB Next Multiplex Small RNA Library Prep Kit (New England Biolabs) was used for library preparation, as directed by the manufacturer, and miRNA sequences were mapped to miRBase 21.

### Analysis of single cell RNA-seq data

Published amacrine cell single cell RNA-seq data (Yan et al., 2020) was downloaded from GEO (GSE149715) and used to generate a gene expression table with normalization and batch correction using the Seurat package (https://satijalab.org/seurat/) as described (Yan et al., 2020). Pearson correlation of Foxn3 expression with the other 18204 genes passing Seurat filtering was performed using R cor.test. See Supplemental Table S5.

### CRISPR Sequencing analysis

Genomic DNA was prepared from GFP-positive retina explants 48 hours after electroporation with Foxn3 CRISPR vectors. Genomic DNA from two pooled explants was purified by using DNeasy Blood & Tissue Kit (Qiagen) for each sample. Q5 high-fidelity DNA polymerase (NEB) was used to PCR amplify genomic DNA near each Cas9 target site. PCR primers are listed in Supplemental Table S6. PCR products were gel purified, then analyzed by Illumina sequencing (Massachusetts General Hospital CCIB DNA Core). Sequence reads were analyzed using CRISPResso (Pinello et al., 2016).

## Acknowledgements

We thank our colleagues who generously provided reagents: Marina Pasca di Magliano and Chris Wright for Ptf1a-Cre mice, Helena Edlund for the anti-Ptf1a antibody, and Mike Uhler for plasmids. We thank Tom Glaser, Brenda Bass, David Chen, Robert Thompson, Anne Vojtek, Beth Rousseau, and Mike Uhler for helpful discussions. Support (DLT): NEI R01 EY024996, NEI R21 EY018707, and a grant from Midwest Eye Banks. This work utilized the Vision Research Core at the University of Michigan Kellogg Eye Center, which is supported by NEI P30 EY007003.

