## Supplemental Figures for "Regulation of retinal amacrine cell generation by miR-216b and Foxn3"

### Supplemental Figures S1-S8

Supplemental Figure S1

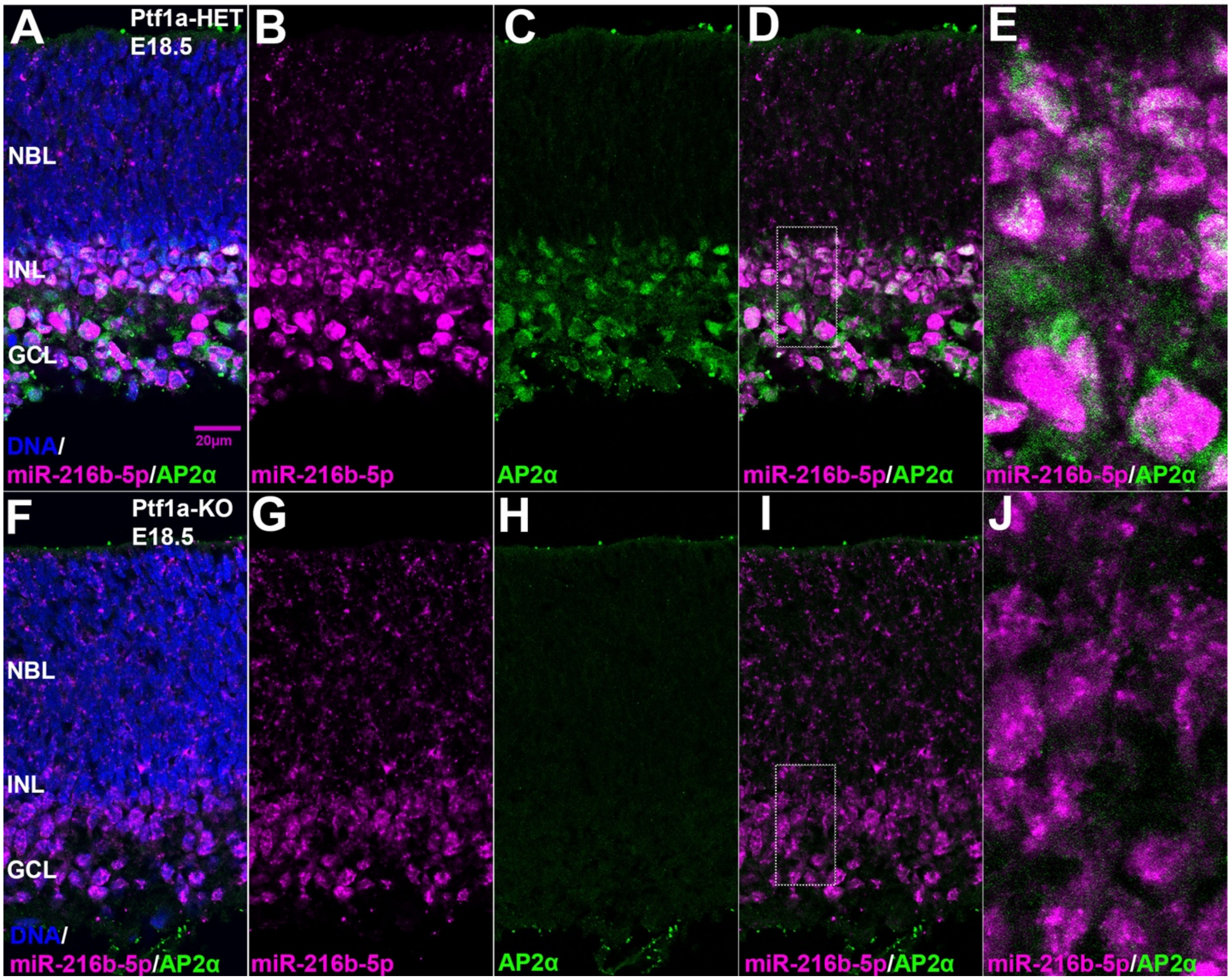


Supplemental Figure S1. miR-216b expression in amacrine cells in E18.5 retinas. (A-E) Mature miR-216b detected by fluorescent in situ hybridization (magenta) overlaps with AP2α (green) in retinas from mice heterozygous for Ptf1a.(F-J) Mature miR-216b is reduced and AP2α is absent in retinas from mice homozygous for disruption of Ptf1a. (E) and (J) show enlargement of regions indicated in (D) or (I). Blue: nuclear DNA.

**Supplemental Figure S2**

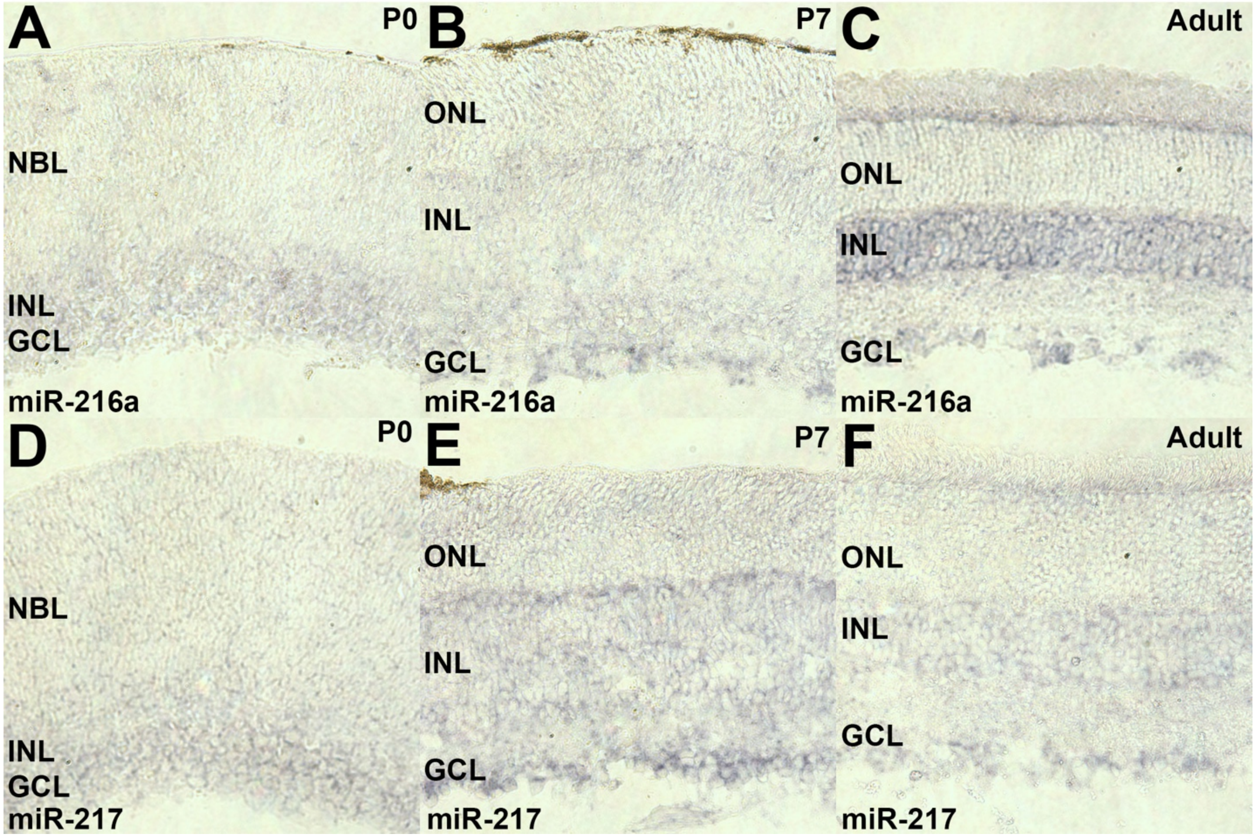


Supplemental Figure S2. miR-216a and miR-217 expression in retina. (A-C) Mature miR-216a detected by in situ hybridization (purple) is present in INL and GCL of P0, P7, and adult retinas. (D-F) Mature miR-217 is present in INL and GCL of P0, P7, and adult retinas.

Supplemental Figure S3

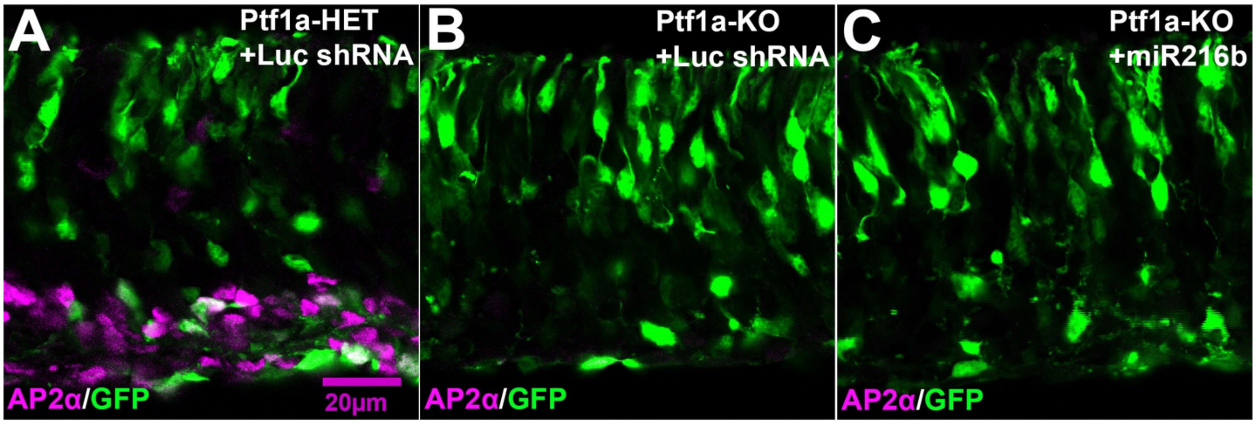


Supplemental Figure S3. Ectopic expression of miR-216b in the developing retina does not generate amacrine cells in retinas from Ptf1a knockout mice. Plasmid miRNA expression vectors which co-express GFP and either pre-miR-216b or the control Luc shRNA were introduced into retinas at E16.5 by electroporation, then the retinas were maintained in explant culture for 8DIV. (A) Control retina from a Ptf1a heterozygous mouse showing AP2α (magenta) overlaps with a subset GFP-labeled cells (green). (B, C) In retina explants from mice homozygous for disruption of Ptf1a, AP2α labeled cells were not present among the GFP-labeled cells (representative images, N=3 for each vector in Ptf1a knockout retinas).

**Supplemental Figure S4**

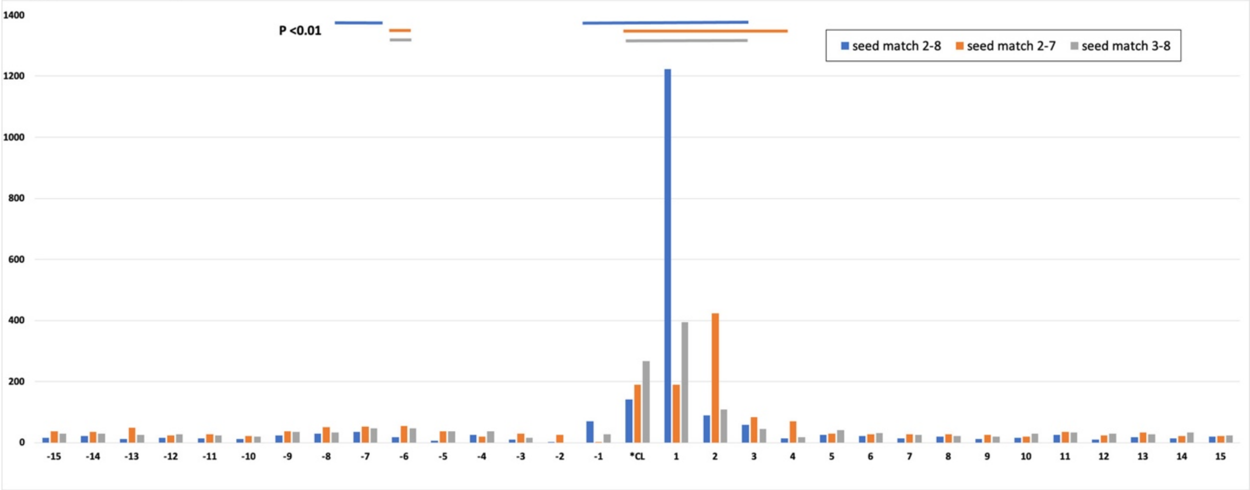


Supplemental Figure S4. miRNA seed matches are enriched near the predominant crosslinks in retina Argonaute PAR-CLIP libraries. We mapped common retinal miRNA seed sequences in 31nt windows around 7237 predominant crosslink sites in 3' UTR exons. Crosslink sites near exon borders were excluded to avoid seed mappings outside of the exons. Seed matches were mapped at the 5' most base of each match. The graph shows the count of 3 classes of seed matches +/-15nt relative to the predominant crosslink site (*CL), summed across all sites. Only the longest match to a seed is counted (i.e. a 2-7 match or a 3-8 match located within a 2-8 match are not counted). Positions with enrichment ($P < 0.01$) for each type of seed match are indicated by matching color lines above the graph. Seed matches which overlap with the crosslink position must have an A at the appropriate position in the seed to basepair with the crosslinked U in the target, so fewer seeds can map at positions -6 to 0. miRNA seeds used: miR-9-5p CTTTGGT, miR-9-3p AAAGCTA, let-7ifagcdbe-5p/miR-98-5p GAGGTAG, miR-124-3p AAGGCAC, miR-182-5p TTGGCAA, miR-183-5p ATGGCAC, miR-26ab-5p TCAAGTA, miR-181abdc-5p ACATTCA, miR-30dcaeb-5p GTAAACA, miR-148ab-3p/miR-152-3p CAGTGCA, miR-25-3p/miR-92ab-3p/miR-32-5p ATTGCAC, miR-99ba-5p/miR-100-5p ACCCGTA, miR-93-5p/miR-20a-5p/miR-17-5p/miR-106b-5p AAAGTGC, miR-125ab-5p/miR-351-5p CCCTGAG, miR-7ab-5p GGAAGAC.

**Supplemental Figure S5**

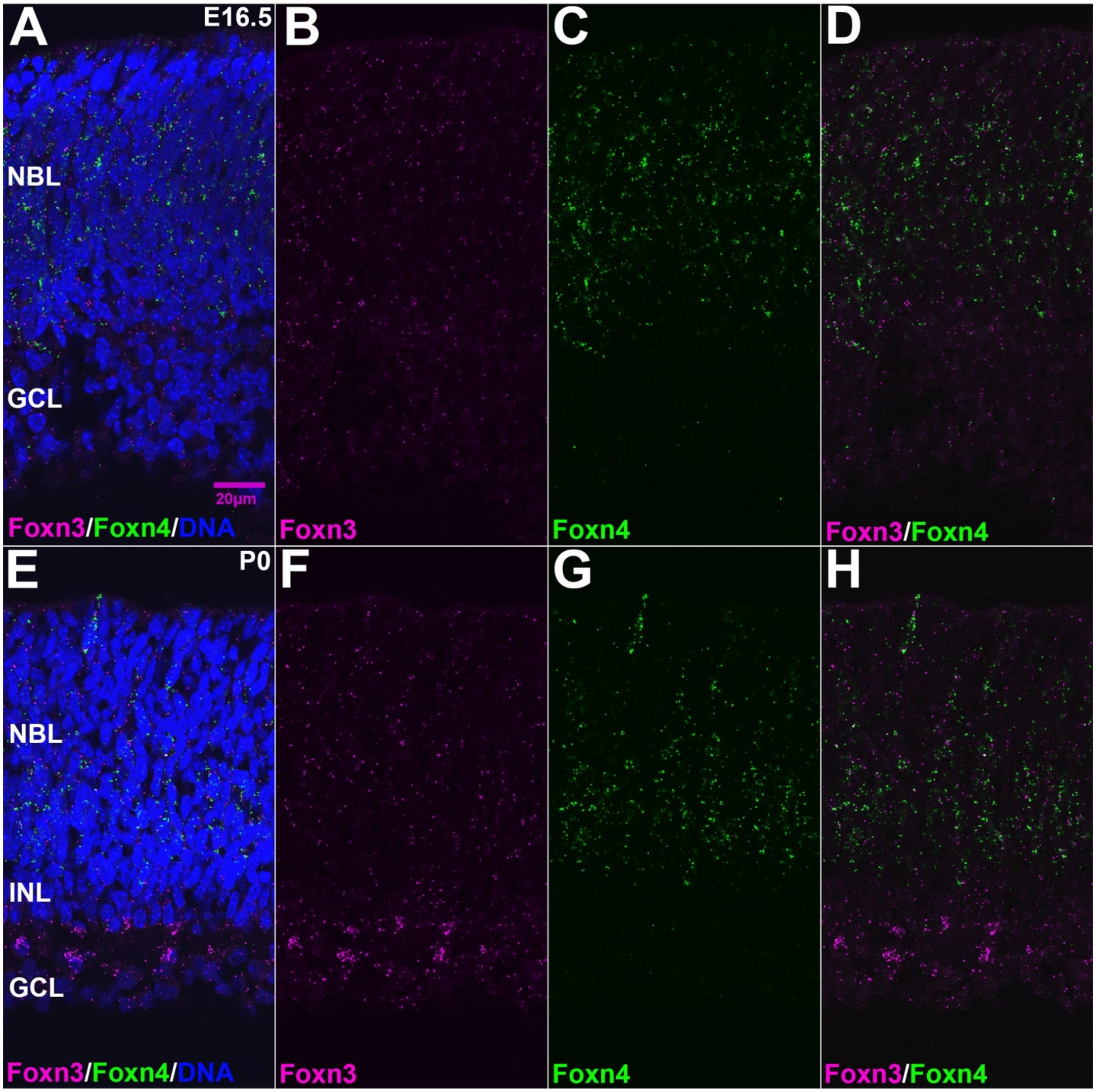


Supplemental Figure S5. Overlap between Foxn3 and Foxn4 expression in the developing retina. (A-D) Foxn3 (magenta) and Foxn4 (green) expression in cells of the E16.5 retina, detected by in situ HCR. (E-H) Complete panels for Foxn3 and Foxn4 expression detected by in situ HCR in the P0 retina; Panel H is the same as Fig. 4G. Blue in A and E: nuclear DNA.

Supplemental Figure S6

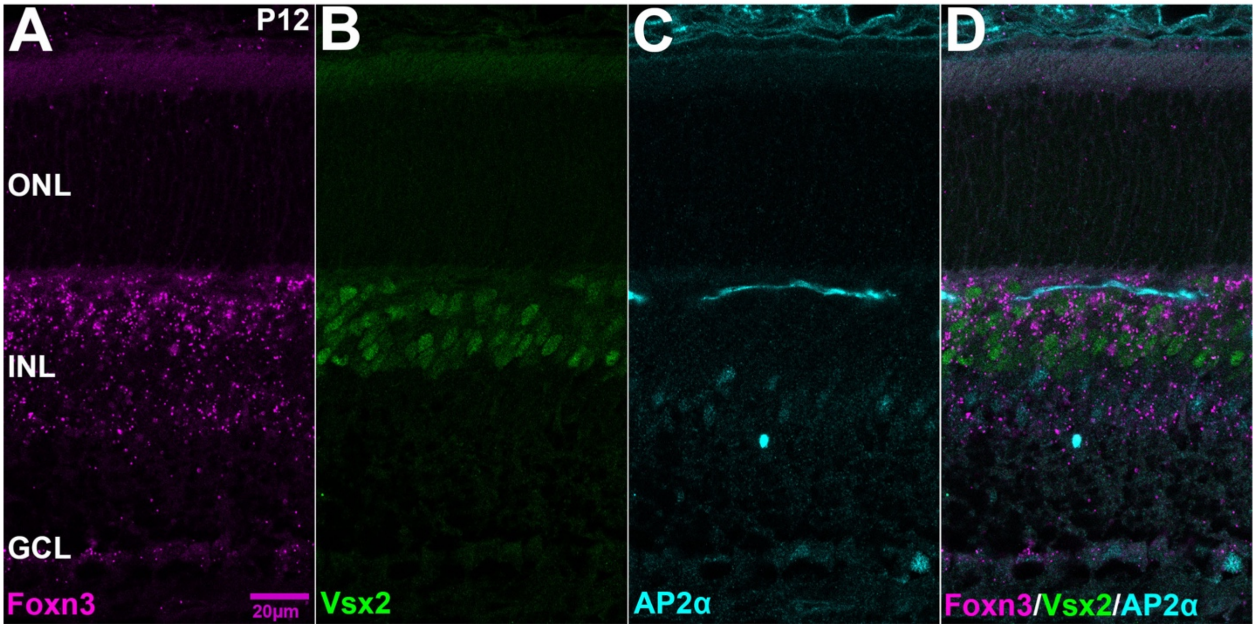


Supplemental Figure S6. Complete panels for Foxn3 and marker expression in the P12 retina.

(A-D) Foxn3 mRNA in the INL at P12 (magenta), detected by in situ HCR, overlaps with Vsx2 (green) and AP2α (cyan) detected by immunofluorescence. Panel D is the same as Fig. 4K.

**Supplemental Figure S7**

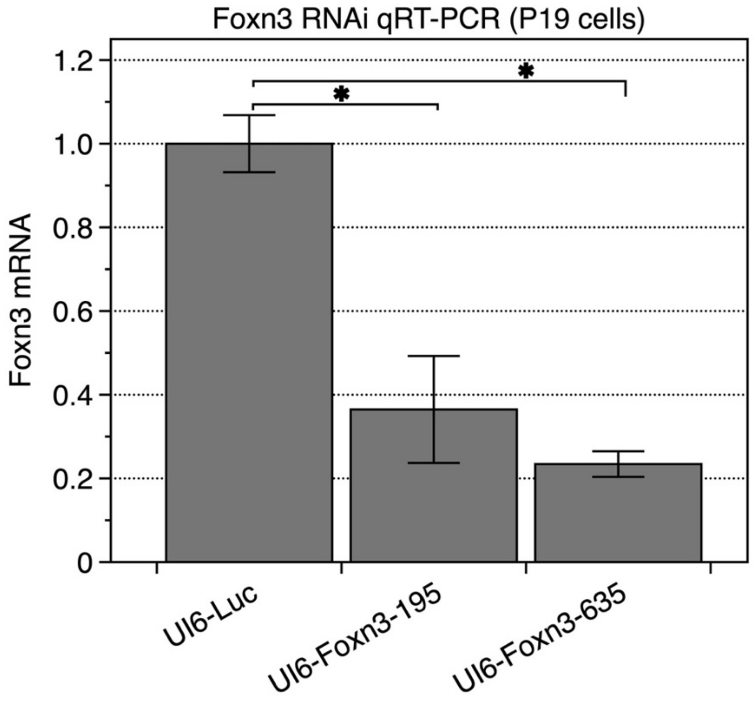


Supplemental Figure S7. Foxn3 RNAi reduces Foxn3 mRNA in a cell line. Mouse P19 cells were transfected with the control Luc-shRNA vector or one of two Foxn3 shRNA vectors. Foxn3 mRNA was measured by qRT-PCR. Both Foxn3 shRNAs reduced endogenous Foxn3 mRNA relative to the control (N=3). * $P < 0.05$.

**Supplemental Figure S8**

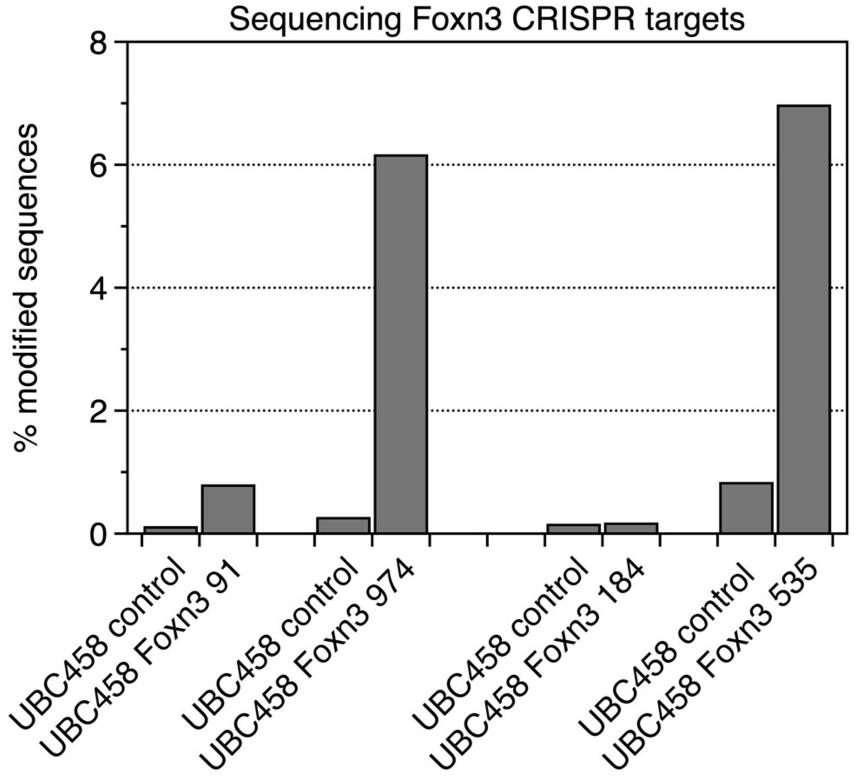


Supplemental Figure S8. Illumina sequencing of genomic PCR products shows disruption of Foxn3. Each Foxn3 CRISPR/Cas9 vector expresses two sgRNAs, targeting different sequences in the Foxn3 coding region. Foxn3 CRISPR1: sgRNA 91 + sgRNA 974; Foxn3 CRISPR2: sgRNA 184 + sgRNA 535. PCR products corresponding to each of the two different targets for each vector were amplified from genomic DNA from pooled GFP-positive retinas electroporated with either a Foxn3 CRISPR vector or the parental pUBC458 Cas9 vector and maintained as retinal explants for 2 DIV. Products were analyzed by Illumina sequencing to count modified sequences (relative to the mm10 reference sequence). For each of the two Foxn3 CRISPR vectors, one of the two sgRNAs is more effective.

**Supplemental Tables (separate Excel files):**

Supplemental Table S1. miRNA counts for small RNA seq of E16.5 retinas from mice heterozygous or homozygous for disruption of Ptf1a.

Supplemental Table S2. miRNA counts for small RNA seq of P0 retinas from wild-type CD-1 mice.

Supplemental Table S3. PAR-CLIP crosslink sites with a nearby sequence matching the seed sequences of both miR-216b and miR-216a.

Supplemental Table S4. PAR-CLIP crosslink sites with a nearby sequence matching a seed sequence of either miR-216b or miR-216a but not both.

Supplemental Table S5. Foxn3 mRNA expression in amacrine single cell RNA-seq data.

Supplemental Table S6. Sequences of oligonucleotide probes and primers.
